# Resource conservation manifests in the genetic code

**DOI:** 10.1101/790345

**Authors:** Liat Shenhav, David Zeevi

## Abstract

Nutrient limitation is a strong selective force, driving competition for resources. However, much is unknown about how selective pressures resulting from nutrient limitation shape microbial coding sequences. Here, we study this ‘resource-driven’ selection using metagenomic and single-cell data of marine microbes, alongside environmental measurements. We show that a significant portion of the selection exerted on microbes is explained by the environment and is strongly associated with nitrogen availability. We further demonstrate that this resource conservation optimization is encoded in the structure of the standard genetic code, providing robustness against mutations that increase carbon and nitrogen incorporation into protein sequences. Overall, we demonstrate that nutrient conservation exerts a significant selective pressure on coding sequences and may have even contributed to the evolution of the genetic code.

## Introduction

Nitrogen and carbon are major limiting factors in many ecosystems, with recent studies linking their availability to core genomic properties (*1, 2*). In low-nitrogen environments there is a lower incorporation of nitrogen-rich side-chains into proteins, a strong A+T bias in nucleotide sequences, and smaller genome sizes (*3–5*). An opposite trend was shown for carbon (*2*) and indeed nitrogen and carbon concentrations are typically inversely correlated (*6*). These studies suggest a purifying selective pressure associated with resource conservation, which we term ‘resource-driven’ selection.

This resource-driven selection postulates that DNA mutations which result in excess incorporation of nutrients such as nitrogen and carbon into proteins, are disfavored. However, not all mutations have the same effect on protein sequences due to constraints imposed by the structure of the genetic code. The standard genetic code, common to virtually all of life on earth, is known to be highly efficient in mitigating deleterious effects of mistranslation errors and point mutations (*7*), specifically those leading to radical changes in amino acids. This error minimization is prominent among theories regarding the origin of the genetic code (*8–11*), which propose that the code had evolved through selection to minimize potential adverse effects of mutations on protein structure and function (*12–14*). To quantify code optimality, some theories provide structurally-informed amino acid metrics based on hydropathy and stereochemistry [e.g., the polar requirement scale (*11*) and hydropathy index (*15*)]. To our knowledge, an optimization of nutrient conservation in the genetic code has not been studied thus far.

Here, we use publicly available metagenomic and single-cell data from global marine habitats, coupled with their corresponding measurements of environmental conditions, to comprehensively characterize and quantify the selective pressure resulting from nutrient limitation. We estimate selection in 746 marine samples (**Fig. S1A**) and quantify its ‘resource-driven’ component, which we show to be a ubiquitous purifying selective force. We further demonstrate that the structure of the genetic code, known to minimize the effects of DNA mutations on the structure and function of proteins, also minimizes nutrient expenditure given a random mutation. We show that this optimization for resource conservation is orthogonal to known mechanisms of error minimization in protein structure and function encoded in the genetic code. This analysis exposes a hitherto unknown facet of the genetic code suggesting resource conservation as an important contributor to its evolution.

## Results

### Widespread purifying selection in the marine environment

To comprehensively characterize how coding sequences of marine microbes are affected by resource availability, we devised a computational pipeline that calculates selection metrics from marine metagenomic samples (**Fig. S1B**). We downloaded 746 samples from Tara oceans (*16*) (n=136), bioGEOTRACES (*17*) (n=480), and the Hawaii Ocean- (HOT; n=68); and Bermuda Atlantic- (BATS; n=62) Time-series (*17*) [**Fig. S1A**; (*18*)]. We aligned these reads to the Ocean Microbiome Reference Gene Catalog (OM-RGC) (*16*), a database of genes from marine environments that is accompanied by functional information. We then searched for single nucleotide polymorphisms (SNPs) in genes that had sufficient high-quality coverage [**Fig. S1B**; (*18*)]. Overall, we found 71,921,864 high-confidence SNPs (*18*), in a total of 1,590,843 genes.

To quantify purifying selection on different gene functions, we annotated genes from the OM-RGC database using either KEGG orthology [KO; (*19*)] or eggNOG orthologous groups [OG; (*18, 20*)]. Using called SNPs, we calculated for each orthologous group and in each sample, the ratio of nonsynonymous polymorphisms which lead to a change in the coded amino acid (pN), to synonymous polymorphisms, which maintain it (pS) [pN/pS; (*18*)]. Across all samples, we found pN/pS ratios to be close to zero with an average of 0.074 in OGs (CI [0.072, 0.075]) and 0.079 in KOs (CI [0.077, 0.080]; **Fig. 1A**), indicating widespread purifying selection in the marine environment. To corroborate the validity of calculating selection metrics from metagenomic samples, we compared nonsynonymous mutations leading to “conservative” amino acid substitutions to those leading to “radical” substitutions and found conservative mutations to be significantly more common (**Supplementary text**).

**Figure 1.**
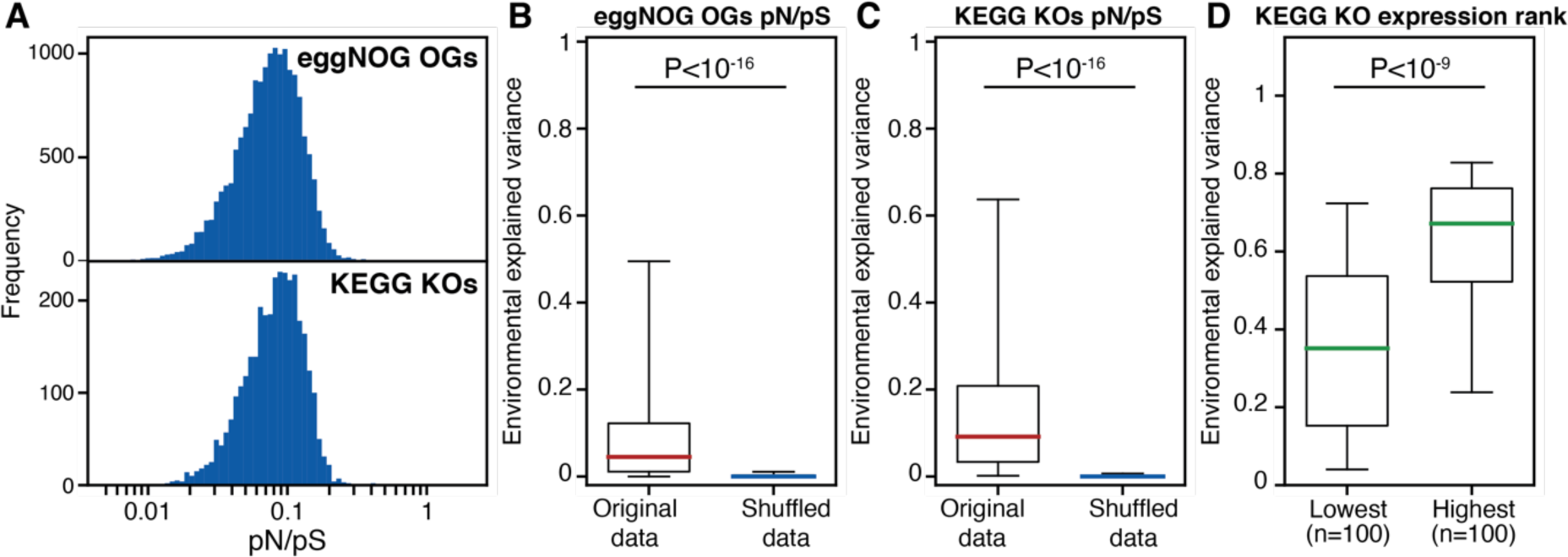
Analysis of pN/pS reveals resource-driven selection. (A) Histogram of pN/pS ratios for eggNOG orthologous groups [OG (20); top] and KEGG orthologs [KO (19); bottom] across all marine samples. (B) Box plot (line, median; box, IQR; whiskers, 5th and 95th percentiles) of the variance of eggNOG OG pN/pS that was explained by the environment in a linear mixed model [LMM;(18)] as compared to the same data with shuffled labels. (C) Same as B, for KEGG KO pN/pS. P, Wilcoxon signed-rank test. (D) Box plot of variance in pN/pS explained by the environment in the 100 lowest and highest expressed KEGG KOs. P, Mann-Whitney U test.

### Significant resource-driven selection across marine microbial genes

Based on recent studies (*2–5*), we hypothesized that nutrient availability is a central driver of this purifying selection. We sought to quantify the environmental contribution to this selective pressure. We considered environmental measurements taken alongside each sample, including depth, water temperature and salinity, and the concentrations of nitrate, nitrite, oxygen, phosphate and silicate [**Fig. S2A-H**; (*18*)].

These environmental measurements presented consistent correlation patterns with the pN/pS of many KEGG and eggNOG orthologs (**Fig. S3**). However, as they are also highly correlated with each other (**Fig. S2I**), we cannot accurately estimate their individual effects. We therefore sought to estimate the fraction of variance in pN/pS explained by the environment while accounting for these covariations. We used a linear mixed model (LMM) with variance components (*18*), which allows to estimate the fraction of variance in pN/pS ratios (dependent variable) that is explained by the environment (random effect) while controlling for the complex correlation structure between the environmental parameters. We term the fraction of variance in pN/pS explained by resource availability ‘environmental explained variance’ [EEV; (*18*)]. Across both KEGG and eggNOG orthologs we found that a significant fraction of the variance in pN/pS can be attributed to the environment (**Fig. 1B,C**, **S4A,B**; Mann-Whitney *U* test P<10^-16^), with nitrate being more strongly correlated with pN/pS ratios than any other environmental parameter (**Fig. S5**; Kolmogorov-Smirnov test; P<10^-30^ for all comparisons). Examining typical DNA mutations and amino acid substitutions in nitrate-rich versus nitrate-poor environments, we find that environmental nitrate is associated with specific changes to both DNA and protein sequences, favoring lower nitrogen incorporation into protein sequences when nitrate is scarce (**Supplementary text**).

This association between environmental measurements and the magnitude of purifying selection is significant even after controlling for potential confounders such as time or effective population size (**Fig. S4C,D**; Mann-Whitney *U* test P<10^-16^; **Supplementary text**) as well as in specific environmental niches (**Fig. S6**; Mann-Whitney *U* test P<10^-20^; **Supplementary text**). Additionally, these results were replicated using benchmarking data of assembled genomes from uncultivated single cells from three dominant lineages of the surface ocean [SAR-11, SAR-86 and Prochlorococcus; **Fig. S4E-G**; **Supplementary text**; (*21*)]. These validations demonstrate that the association between selective pressure and environmental conditions is robust to both data type and selection metric and is not confounded by population properties and clade-specific metabolism.

### Environmental association is stronger in resource-consuming genes

With nitrate being the environmental factor most strongly associated with pN/pS, we sought to provide a population-level perspective to the theory and experiments demonstrating selection against mutations that increase the nitrogen requirements of cells, especially in nitrogen-limited conditions (*3, 5, 22*). This implies stronger purifying selection in highly expressed genes (*3, 4*), where one DNA mutation could translate to thousands of proteins, each consuming more resources (illustrated in **Fig. S7**). We used an additional expression dataset for marine microbial genes (*23*) to rank KEGG orthologs by their mean expression (*18*) and found that the 100 most highly expressed KEGG KOs had a significantly higher EEV as compared to the 100 least expressed ones (**Fig. 1D, S8**; Mann-Whitney *U* test P<10^-9^). We replicated these results using single-cell data pertaining to specific bacterioplankton lineages (Mann-Whitney *U* test P<10^-7^). Additionally, we found that extracellular protein-coding genes, where resources excreted from the cell cannot be recycled, had significantly higher EEV as compared to other gene groups [**Fig. S9**; P<0.05; (*18*)]. This higher EEV in resource-consuming genes further strengthens our results regarding the breadth of resource-driven selection.

### Resource-conservation as an optimization mechanism in the genetic code

We observed selection against DNA mutations which result in excess incorporation of nutrients, such as nitrogen and carbon, into proteins. However, not all mutations have the same effect on protein sequences due to constraints imposed by the structure of the genetic code. Indeed, the genetic code was shown to minimize the impact of point mutations on protein structure and function (*12–14*). We hypothesized that the genetic code also minimizes the impact of point mutations on nutrient incorporation into proteins. Specifically, that the genetic code acts as a buffer between the DNA level, where mutations occur, and the protein level, where resource-driven selection is exerted, such that the expected cost of a random DNA mutation, in terms of added resources to proteins, is minimized.

To rigorously test this hypothesis, we first defined, for each element *e* (e.g., carbon, nitrogen), a function quantifying the cost of a single mutation as the added number of amino acid atoms resulting from it. For example, a missense mutation from codon CCA to CGA results in an amino acid substitution from proline to arginine, with an increase of one carbon and three nitrogen atoms, setting the nitrogen cost of such mutation to 3 and the carbon cost to 1 (**Fig. 2A**).

**Figure 2.**
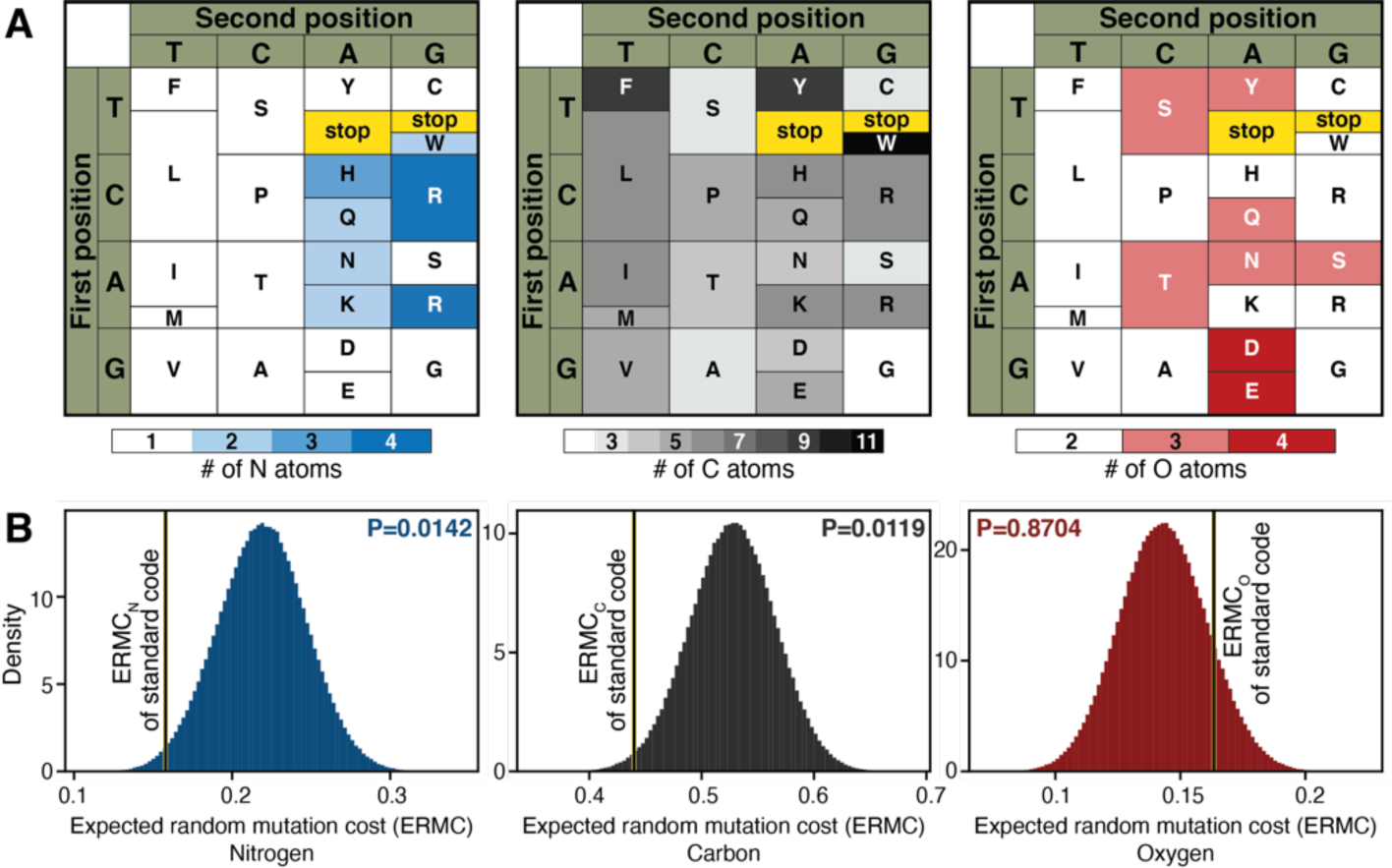
Resource-conservation is facilitated by the genetic code. (A) Nitrogen (left), carbon (center) and oxygen (right) content of different amino acids depicted along their positions in the standard genetic code. (B) Histograms of the expected random mutation cost (ERMC), in 1,000,000 random permutations of the genetic code for nitrogen (left, blue), carbon (center, black) and oxygen (right, red). Black-yellow bar marks the ERMC of the standard genetic code, ERMC(Vs), for each of the elements.

To estimate the cost of random mutations across the entire genetic code, we calculated, per element (e.g., nitrogen), the Expected Random Mutation Cost (ERMC):

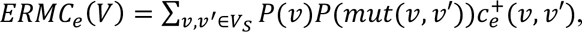

Where V is a genetic code; P(v) the abundance of codon v, here calculated from all marine samples; P(mut(v,v′)) the probability of mutation from codon v to codon v’, set to be the relative abundance of the single nucleotide mutation driving this codon change (e.g., for mutation from GCA to CCA, we use the abundance of G-to-C transversions); and the cost function c_e_^+^(v,v′) the number of atoms of element *e* added post-mutation (*18*). For the standard genetic code, and codon abundances and mutation rates calculated from marine microbes, we report an ERMC of 0.440, 0.158 and 0.163 for carbon, nitrogen and oxygen, respectively, corresponding to an average increase of this number of atoms per random mutation (**Fig. 2B**).

To check if the standard genetic code, along with codon abundances and mutation rates, is indeed robust to resource-consuming mutations, we compared it to other hypothetical codes. We simulated alternative genetic codes by randomizing the first and second position in all codons one million times, to create a null distribution of ERMC (*18*). We found that the standard genetic code, common to most life forms, is parsimonious in terms of carbon and nitrogen utilization, given a random mutation. This is marked by a significantly low ERMC for nitrogen (**Fig. 2B**; ERMC_N_ P=0.0142) and carbon (ERMC_C_ P=0.0119), but not oxygen (ERMC_O_ P=0.8704).

We compared the extent of robustness against an addition of carbon or nitrogen to well-studied structure and function error minimization mechanisms (*12–14*). We calculated ERMC for changes in hydropathy (*15*) and polar requirement [PR; (*11*)], both structurally-informed amino acid properties used to determine code error minimization (*18*). We found that these optimization mechanisms are of a similar magnitude to those shown here for nitrogen and carbon [**Fig. S10C,D**; ERMC_hyd_ P=0.0152; ERMC_PR_ P=0.0137; (*18*)].

We next sought to confirm that optimization for carbon and nitrogen is not confounded by these known optimization mechanisms. To this end, we devised a hierarchical model which examined the subset of genetic codes (out of 1M hypothetical codes tested) that have a lower ERMC than the standard code for PR or hydropathy, and tested whether this subset is also optimized for nitrogen or carbon (*18*). If nutrient optimization exists separately from structural optimization, we expect the standard code to be optimized for carbon and nitrogen even as compared to this subset, featuring significantly lower ERMC_N_ and ERMC_C_. We find that out of 15,223 hypothetical codes that have ERMC_hyd_ lower than the standard code, only 270 have lower ERMC_N_ (**Fig. 3, S10E**; P=0.019) and only 249 have lower ERMC_C_ (**Fig. 3, S10E**; P=0.0208). Similarly, out of the 13,729 hypothetical codes that have ERMC_PR_ lower than the standard code, only 83 have lower ERMC_N_ (**Fig. S10F**; P=0.0058) and only 442 have lower ERMC_C_ (**Fig. S10F**; P=0.037). This is in contrast with the observed overlap between hydropathy and PR: out of the 15,223 hypothetical codes that have ERMC_hyd_ lower than the standard code, 6,736 have lower ERMC_PR_ (**Fig. S10E**; P=0.4425). These results indicate that the detected carbon and nitrogen optimization is not confounded by previously reported optimization properties such as hydropathy and PR.

**Figure 3.**
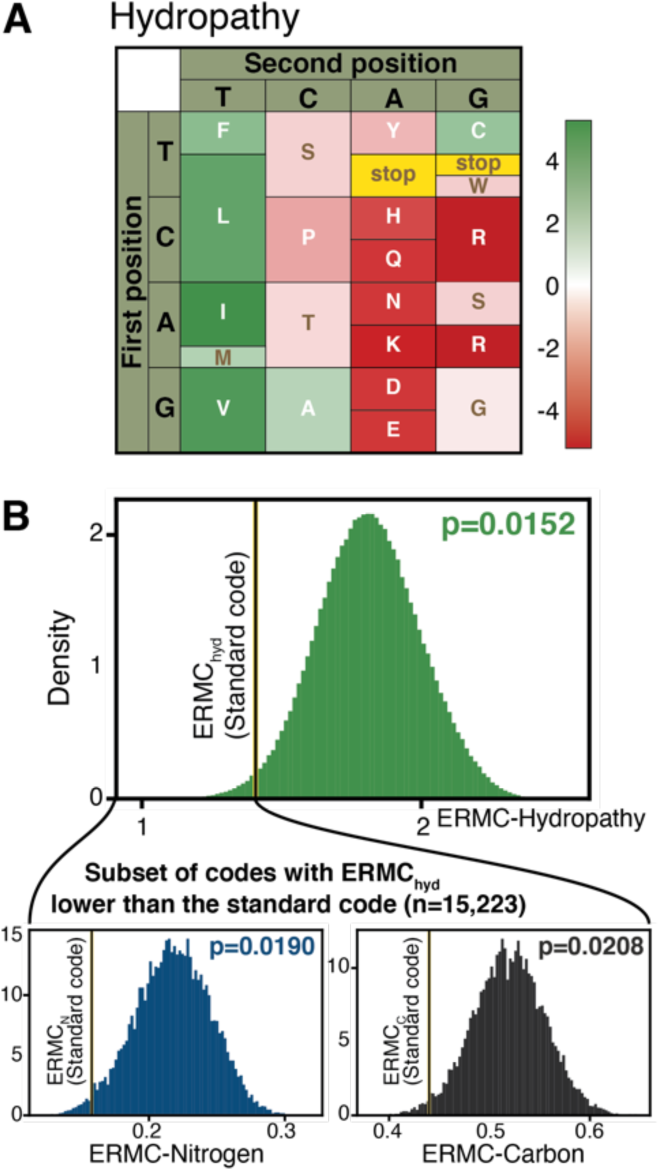
Optimization for carbon and nitrogen is not confounded by hydropathy. (A) Hydropathy of different amino acids depicted along their positions in the standard genetic code. (B; top) As in Fig. 2B, for hydropathy; (B; bottom) histograms of ERMC for nitrogen (left) and carbon (right) for the subset of hypothetical genetic codes with ERMC lower than the standard code for hydropathy. P, permutation test.

Remarkably, only 128 out of 1M randomized genetic codes were better than the standard code in conservation of nitrogen and carbon together (**Fig. S10G,H**; ERMC_CN_ P=1.28x10^-4^). We note that this number is significantly smaller than the number of hypothetical codes expected to have both a lower ERMC_C_ and ERMC_N_ (Chi-square test of independence P=0.0013; **Table S1**). This is possibly driven by a small overlap between the positions of high-nitrogen and high-carbon amino acids. This property of the standard code potentially enables concurrent optimization for both carbon and nitrogen. These results highlight a new optimization principle of the genetic code that is of similar magnitude- and independent of- previously proposed principles.

### The genetic code facilitates resource conservation across kingdoms

To show that the resource-robustness of the genetic code was not limited to our dataset and analytic approach, we calculated the ERMC of 187 strains of genera *Prochlorococcus* and *Synechococcus*. We computed codon abundances and mutation rates using published protein-coding sequences (*1*) and the accepted transition:transversion rate of 2:1 (*18, 24*). We corroborated significant conservation of carbon, nitrogen, and both elements combined (**Fig. S11A**; ERMC_C_ mean P=0.013, P=0.020; ERMC_N_ mean P=0.049, P=0.032; ERMC_CN_ P=0.0004, P=0.0007 for *Prochlorococcus* and *Synechococcus*, respectively).

To explore whether this nutrient conservation optimization mechanism of the standard code extends across organisms, we performed a similar calculation using codon abundances from 39 organisms across all domains of life, including all human protein-coding sequences, and a range of transition:transversion rates (*18*). Similarly to marine microbes, we find that the standard code is optimized in terms of resource utilization for all tested organisms, marked by a significantly low ERMC for nitrogen and carbon combined, across all transition:transversion rates (P<0.01, **Fig. 4**). Moreover, we find significant optimization even in the theoretical case where all codon abundances are the same (**Fig. S11B**). The codon abundances of a great majority of organisms also demonstrate significantly low ERMC for nitrogen (**Fig. S11C**) and carbon (**Fig. S11D**), for a wide range of transition:transversion rates. These results indicate that resource optimization in the genetic code transcends taxonomy, codon choices, and mutation rates.

**Figure 4.**
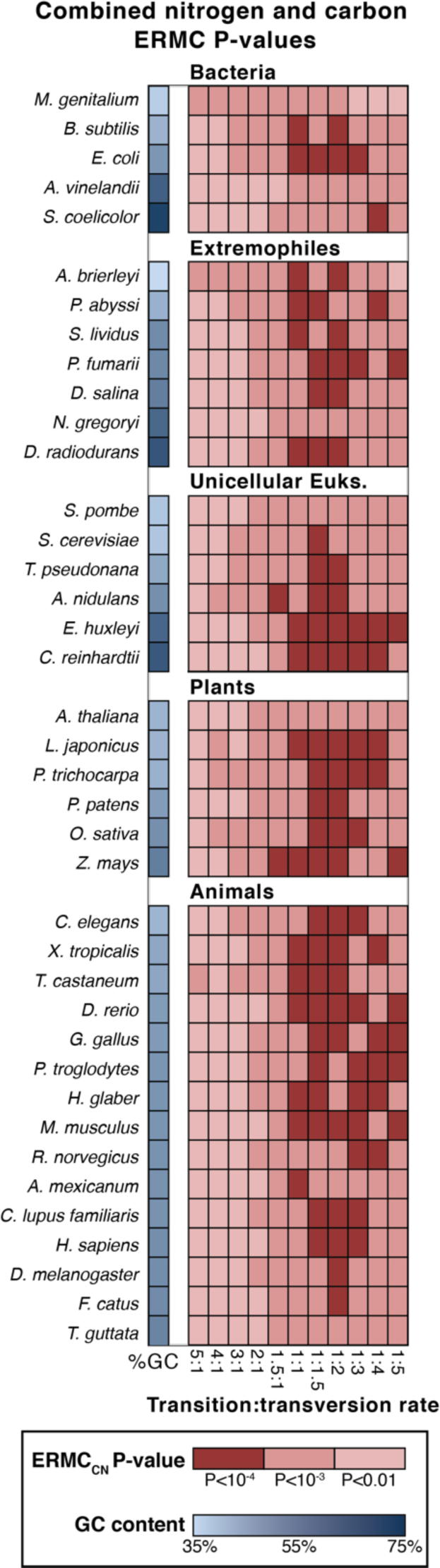
The genetic code is optimized for resource conservation across organisms. Heat map of ERMC_CN_ P-values across 39 organisms and 11 transition:transversion rates. Organisms in each group are ordered by the GC content of their coding sequences.

### Resource conservation may bias codon usage

We hypothesized that variation in the third codon position for a single amino acid and may also be biased due to the differential cost of a random mutation for each codon. We therefore focused on codon usage of the amino acid threonine. We note that a C-to-G transversion in the second position for codons ACT and ACC yields serine (AGT and AGC, respectively), but the same mutation for codons ACA and ACG yields arginine (AGA and AGG; **Fig. 5A**, inset). Arginine has higher carbon and nitrogen content than serine. We thus hypothesized that for a cell to conserve nutrients in case of a random mutation, codons ACT and ACC should have a higher abundance than codons ACA and ACG, respectively, given a known GC-bias that is a genomic quantity. To test this hypothesis, we examine codon usage in 187 *Prochlorococcus* and *Synechococcus* strains, and show a significantly higher use of ACT as compared to ACA (**Fig. 5A**; Wilcoxon signed-rank test P<10^-20^) and ACC as compared to ACG (**Fig. 5A**; Wilcoxon signed-rank test P<10^-20^). Similarly, isoleucine codon ATT has higher abundance as compared to ATA (**Fig. 5A**; Wilcoxon signed-rank test P<10^-20^). These results demonstrate that resource conservation may guide codon usage and thereby affect not only protein sequence but also cellular translation efficiency.

**Figure 5.**
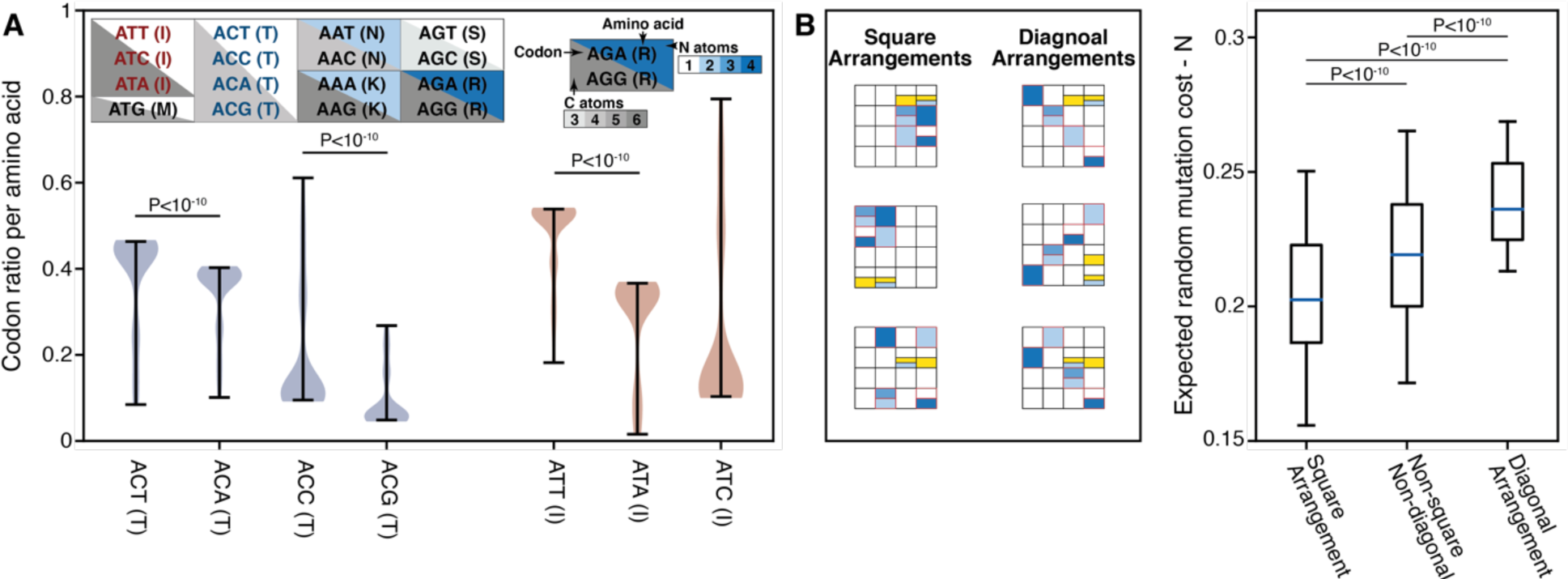
Structural properties and codon usage bias underlying optimality in the genetic code. (A) Violin plot of codon usage among 187 species of Prochlorococcus and Synechococcus showing significant preference of threonine codons ACT and ACC as compared to ACA and ACG, and of isoleucine codon ATT as compared to ACA. P, Wilcoxon signed-rank test. (B) Box plot (line, median; box, IQR; whiskers, 5th and 95th percentiles) of ERMCN of square arrangements (left) and diagonal arrangements [right, (18)], as compared to all other arrangements (center) out of 10,000 hypothetical arrangements. P, Mann-Whitney U test.

### Structural principles drive optimization in the genetic code

We next examined the organizing principles underlying code resource optimization. We observed that codons of the nitrogen-rich amino acids histidine, glutamine, asparagine, lysine and arginine span only two nucleotides in their first position and two in their second position. We define this organization to be a ‘square’ arrangement, and hypothesize that it amplifies nitrogen conservation [**Fig. 5B**; (*18*)]. Specifically, in the square arrangement, some codons require at least two mutations to increase the number of nitrogen atoms (e.g., those coding for alanine and valine). This is in contrast to other hypothetical arrangements, including a ‘diagonal’ one in which nitrogen-rich amino acid codons span all possible nucleotides in the first and second positions [**Fig. 5B**; (*18*)]. We suggest that this arrangement would be nutrient-wasteful, as in such arrangements a single mutation in either the first or second codon position could increase the nitrogen content of a protein. To test this, we generated 10,000 hypothetical codes, with 220 arrangements happening to embody a square structure, and 127 a diagonal one. We found that, when compared to all other possible arrangements, square arrangements present a significantly lower ERMC_N_ while diagonal arrangements exhibit a significantly higher ERMC_N_ (**Fig. 5B**; Mann-Whitney *U* test P<10^-10^ for both). This result demonstrates that resource optimization in the standard code is driven by structural principles, perhaps underlying the optimization observed across kingdoms.

## Discussion

In this work, we characterize and quantify the selective forces exerted by nutrient availability on protein coding genes in marine environments. We provide a data-driven, population-level perspective and show that resource-driven selection is a ubiquitous force. We further show that a significant portion of DNA mutations may not result in increased nutrient incorporation into protein sequences, due to the structure of the standard genetic code. We provide evidence that the genetic code, shared among most lifeforms, along with organismal codon frequencies, is robust against the incorporation of additional atoms of nitrogen and carbon given a random mutation. We quantitatively demonstrate that this nutrient conservation is not confounded by known error minimization mechanisms of the genetic code.

In light of these results, we hypothesize that resource-driven selection is equally exerted on all parts of the protein-coding gene. This sets it apart from selection to maintain the structural integrity of a protein or the function of its active site, which occur predominantly in structurally important regions (*25*). Thus, accounting for resource-driven selection may improve the identification of alternative translation start sites, alternatively spliced introns, or readthrough stop codons, as intermittently translated regions of the protein may be under weaker resource-driven selection as compared to constitutively translated ones (*26*).

We note that in addition to resource-driven selection there are other forces associated with the environment, driving both selective and neutral mechanisms. Genomic GC-content is associated with exposure to UV radiation (*27*) and nitrogen availability (*2, 28*) yet nonsynonymous mutations, especially in codons encoding structurally and functionally important protein positions, are likely to be less affected by GC-bias. Furthermore, codon usage bias was also shown to be associated with the environment (*29*).

We show that the genetic code optimizes nutrient conservation, calling upon an examination of code evolution theories. Hypotheses regarding the origin of the genetic code include stereochemical affinity between a codon or anticodon and their amino acid (*11, 30*); a frozen accident theory (*31*); a co-evolution of the genetic code with the emergence of amino acid-producing biosynthetic pathways (*9*); and an early fixation of an optimal genetic code, suggesting selection for error minimization (*10, 32*). Our results are in line with the latter theory and suggest that the genetic code may have also been optimized for nutrient conservation. This implies that the genetic code can be viewed as a buffer between the evolutionary forces of mutation and selection, the former occurring in DNA sequences and the latter predominantly in proteins. In the case of nutrient conservation, many DNA mutations do not result in the incorporation of additional nutrients into proteins, and are thus not selected against, which may allow more ‘freedom’ to explore the mutation-space. Notwithstanding, while optimized for resource conservation, the genetic code is also near-immutable. This is evident in heterotrophic eukaryotes, which while being net-nitrogen producers, still harbor a nitrogen-conservative code.

Since the genetic code is common to virtually all organisms, we can assume that it was already fixed in its current form, or one very similar, in the last universal common ancestor. Thus, if the genetic code evolved to optimize resource conservation, it may reflect selective pressures in the beginning of life on earth. An organism harboring a nitrogen- and carbon-efficient genetic code would have had a selective advantage over its peers, especially in the absence of fully evolved DNA mutation repair mechanisms.

## Acknowledgements

We thank Tal Korem and Nimrod D. Rubinstein for discussions, suggestions and help in analyses. We thank Jean-Pierre Eckmann, Otto X. Cordero, Itzhak Mizrahi and Niv Antonovsky for their suggestions.

## Authors contributions

LS and DZ conceived the project, performed analyses and wrote this manuscript.

## Data and materials availability

All data is available in the manuscript, supplementary materials and referenced datasets. Code is available at: https://github.com/zeevilab/resource-conservation

## Supplementary Materials

### Materials and Methods

#### Marine microbiome samples

Marine samples collected with Tara oceans (*16*), bioGEOTRACES (*17*), the Hawaii Ocean Timeseries (HOT) and the Bermuda Atlantic Timeseries (BATS) (*17*) were downloaded from ENA with accessions ENA:PRJEB1787 (TARA oceans prokaryotic fraction), ENA:PRJNA385854 (bioGEOTRACES) and ENA:PRJNA385855 (HOT/BATS), each sample with a minimum of 5 million reads. For all samples for which a size parameter was available, we selected the 0.22µm+ size category (typically between 0.22 and 5µm). This includes TARA ocean samples (prokaryote size) and data pertaining to expression (*22*). In samples from the bioGEOTRACES, HOT and BATS, we used the available size fraction, larger than 0.2µm.

#### Mapping of Illumina reads to reference gene sequences

Samples were mapped to nucleotide sequences from the Ocean Microbiome Reference Gene Catalog (OM-RGC) (*16*) using bowtie2 with parameters --sensitive -a 20 --quiet -p 8 and saved as a bam file using the ‘samtools view’ command. As gene sequences are relatively short, reads from both ends of the metagenomic sequencing samples were mapped separately, and reunited prior to variant calling.

#### Determining metagenomic read assignment probability

We determined the probability of assignment of metagenomic reads to marine microbial genes using the Iterative Coverage-based Read-assignment Algorithm (ICRA) (*32*) with parameters max_mismatch=12, consider_lengths=True, epsilon=1e-6, max_iterations=30, min_bins=4, max_bins=100, min_reads=10, dense_region_coverage=60, length_minimum=300, length_maximum=2e5, use_theta=False. To prevent spurious mapping, alignments were considered for downstream analysis only if the probability of alignment was higher than 0.9.

#### Variant calling

Alignments from both ends of all sequencing runs pertaining to the same sample were united using the samtools cat command and sorted using the samtools sort command, with default parameters.

To facilitate variant calling in tractable timescales, each filtered, untied and sorted bam file was split into chunks, each encompassing 10,000 reference sequences (out of about 40 million reference sequences). For each such batch of reference sequences, we called variants across all samples using the following command: bcftools mpileup --threads 4 -a FORMAT/AD -Q 15 -L 1000 -d 100000 -m 2 -f <OM-RGC fasta> <bam filenames>| bcftools call --threads 4 -Ov -mv -o <output vcf>, where <OM-RGC fasta> is the fasta file of OM-RGC nucleotide sequences, <bam filenames>are the filenames of all bam files pertaining to the reference sequence chunk in question, and <output vcf> is the output vcf file pertaining to that same chunk.

To ensure that sequencing errors do not skew our results, we considered single nucleotide variants as SNPs for downstream analysis only if they had a GATK quality score of at least 30, and an allele frequency of at least 1% (*33*). While the GATK quality requirements filter out sequencing errors randomly distributed across the genes, the 1% criterion eliminates random sequencing errors that accumulate in the same position when depth of coverage reaches very high numbers.

For a sample mapped to a reference gene to be considered for downstream analysis, we demanded that at least 60% of SNPs called along the length of the reference gene for that sample would be supported by at least 4 reads, thereby enabling accurate calculation of pN/pS ratios. For a gene to be considered for downstream analysis, we demanded for that gene to have at least one SNP common to 20 or more samples.

#### Calculation of pN/pS in single genes

While comparing SNP patterns across samples, it is instrumental to avoid biases due to differences in coverage. We therefore downsampled the read coverage (depth) to the minimum across all samples, for each position in a sample that was supported by more than 4 reads. For positions that had minimum support of fewer than 4 reads, no subsampling was performed. Subsampling was performed by drawing from a multinomial distribution, with *n* trials and variant probabilities *p*, where *n* was set to the calculated minimum depth and *p* set to the relative abundance of each variant in a given sample.

The ratio of non-synonymous and synonymous polymorphisms was calculated by considering all called SNPs in every gene. First, we calculate a consensus sequence for each gene by taking, for each SNP position, the variant that was overall more common across all samples (after the subsampling performed above). We counted, for each gene, the number of non-synonymous and synonymous sites across the consensus sequence. For each SNP position in each sample, we counted the number of synonymous and nonsynonymous substitutions. As more than one variant can exist in a single sample, we considered the relative abundance of synonymous to nonsynonymous substitutions dictated by the different variants. For example, if the reference codon was CAC, coding for histidine, one variant, a C-to-G transversion in the third position abundant at 50%, led to a nonsynonymous mutation that resulted in glutamine (CAG) while another variant in the same sample and the same position was a synonymous C-to-T transition, we counted 0.5 synonymous substitutions and 0.5 nonsynonymous substitutions. We followed by calculating the pN/pS ratio:

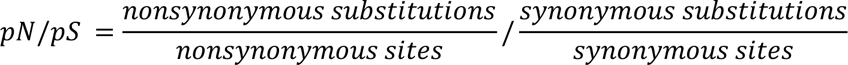

#### Aggregation of calculated metrics using KEGG and eggNOG orthologies

Functional assignments to KEGG KOs and eggNOG OGs for all OM-RGC genes were computed using eggNOG-mapper v2 based on the eggNOG v5.0 database (*19, 34*). For each functional assignment in each sample, all OM-RGC genes annotated with the same functional assignment were concatenated and treated as one long genomic sequence per the calculation of pN/pS ratios. To reduce noise in pN/pS calculation, we considered only KOs and OGs that had at least 5 genes per sample, in at least 50 samples.

#### Calculating nonsynonymous mutations leading to radical and conservative amino acid replacements

We calculated the ratio of nonsynonymous mutations leading to radical amino acid replacements as compared to those leading to conservative replacements using two known amino acid properties: (1) the polar requirement scale (PR) and (2) amino-acid hydropathy index, and multiple thresholds of “radical” and “conservative” (**Table S2**). For each KEGG ortholog and each radical/conservative threshold, we compared the proportion of radical nonsynonymous substitutions (out of all possible radical sites) to the proportion of conservative ones (out of all conservative sites) across all samples using a Mann-Whitney *U* test. P-values were adjusted to control for multiple hypotheses.

Formally, our statistical analysis postulates:

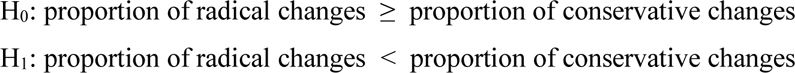

For each radical/conservative threshold, we calculated the proportion of KEGG orthologs in which we reject this null hypothesis and conducted a permutation test. We permuted the sample labels of each radical/conservative threshold across 10,000 iterations, where in each iteration we calculated the proportion of KEGG orthologs in which we reject the null hypothesis. The P-value, per radical/conservative threshold, is defined to be the proportion of iterations in which the ratio of KEGG orthologs in which we reject the null hypothesis is greater than in the original data.

To examine whether highly expressed genes have fewer radical amino acid substitutions as compared to lowly expressed genes, we used data from Pachiadaki et al. (*22*) to rank KEGG orthologs by their mean expression.

#### Benchmarking data of assembled genomes

Benchmarking data was employed in the form of uncultivated single cells of dominant and representative lineages of the surface ocean (SAR-11, SAR-86 and Prochlorococcus) from different parts of the world. We used genomes varying at the strain level (*20*), with potentially minimal batch effects, as those have been analyzed in a single laboratory.

#### Calculation of dN/dS in single genes

For every gene ortholog in each one of the lineages dN/dS was calculated as follows:

1. We extracted all gene sequences pertaining to each ortholog from all strains within each clade. We determined orthologous genes using the annotation that was published alongside the single-cell data. For each clade, we chose the annotation type that had the highest coverage of genes: prokka (*35*) for Prochlorococcus, and Swissprot (*36*) for SAR-11 and SAR-86.
2. To account for the possibility that gene sequences may be misannotated resulting in the grouping of more than one orthologous group under the same name, we performed additional curation by clustering genes using Ward hierarchical clustering (*37*) on protein sequences using normalized edit distance (Levenshtein) as a dissimilarity metric. We cut off clusters that had dissimilarity lower than 30%, to ensure structural similarity.
3. For each of the resulting clusters that had more than 20 members (i.e., genes corresponding to strains within the clade) we generated a multiple sequence alignment (MSA) of protein sequences using MAFFT (*38*), with the following command: mafft --quiet --maxiterate 1000 --localpair <input_fasta> > <output_msa> Following that, we computed a codon alignment using the generated MSA and nucleotide sequences, with the biopython AlignIO package.
4. Using the MSA, we computed a phylogenetic tree of the sequences using raxml (*39*) (raxmlHPC command line version) with model GTRGAMMA; rooted it using the -f I modifier; and computed the ancestral sequence using the -f A modifier.
5. For each leaf node (strain ortholog) in the phylogenetic tree of gene sequences, we calculated dN/dS with respect to the tree root, using the biopython cal_dn_ds function with the “ML” model (Goldman and Yang 1994)(*40*) and default parameters.
6. To quantify purifying selection on different gene functions, we annotated genes from the OM-RGC database using KEGG orthology [KO; (*18*)] and aggregated the dN/dS across genes from the same KO using a geometric mean.

#### Environmental variables

For each sample, we compiled measurements pertaining to the following environmental parameters: Depth [m], Nitrate [µmol/kg], Nitrate+Nitrite [µmol/kg], Oxygen [µmol/kg], Phosphate [µmol/kg], Silicate [µmol/kg], Temperature [°C] and Salinity [g/kg].

Tara oceans metadata was downloaded from PANGAEA (https://doi.pangaea.de/10.1594/) with accession numbers PANGAEA.875575 (Nutrients) and PANGAEA.875576 (Watercolumn sensor), and recorded median values for all the above nutrients were extracted. Tara nutrient concentrations were given as [µmol/l]. Conversion to [µmol/kg] was done by dividing the measured concentration by the measured specific gravity for the same sample.

bioGEOTRACES metadata was compiled from CTD sensor data and discrete sample data from the GEOTRACES intermediate data product v.2 (*41*). HOT metadata was downloaded from the ftp server of the University of Hawai’i at Manoa (ftp://ftp.soest.hawaii.edu/hot/) and BATS metadata was downloaded from the Bermuda Institute of Ocean Sciences (http://bats.bios.edu/bats-data/). As GEOTRACES/HOT/BATS ocean water samples are not linked to specific biological samples as is the case with Tara oceans samples, we considered only water samples from the exact same geographic location, within a day from biological sample collection time, and within 5% difference in depth of collection, and chose the closest sample in terms of time and depth of collection. As all these measurements of environmental conditions are highly correlated with each other (**Fig. S2I**), we utilized this correlation structure to impute missing values using the EM algorithm (*42*).

#### Linear mixed models

We used a linear mixed model (LMM) with variance components, commonly used in population genetics (*43*), which allows to estimate the fraction of variance in pN/pS ratios (dependent variable) that is explained by the environment (random effect) while controlling for the complex correlation structure between the environmental parameters, in a data-driven manner.

#### Generative model

Consider a collection of *M_i_, where i* ∈ {1,2} features (i.e., *M*_1,2_ number of KEGG and eggNOG orthologs respectively), each measured across *N* samples. We get as input an (*M_i_* × *N*) matrix *O^i^*, where 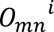 is the pN/pS of ortholog *m* in sample *n*. Let 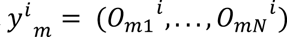 be a *N* × 1 vector representing the pN/pS in ortholog *m*, according to grouping *i*, across *N* samples (e.g., pN/pS in KO K02274 across *N* samples). Let *W* be a (*N* × *q*) normalized matrix of environmental measurements.

This included the depth of the sample, water temperature and salinity, as well as concentration of the key molecules nitrate, nitrite, oxygen, phosphate and silicate.

With these notations, we assume the following generative linear model:

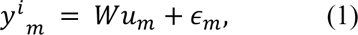

where *u_m_* and *ϵ_m_* are independent random variables distributed as *u_m_* ∼ *N*(0, *σ*^2^ *u_m_^I^*) and 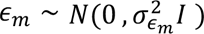. The parameters of the model are *σ*^2^*u_m_* and 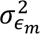.

An equivalent mathematical representation of model (1) is given by

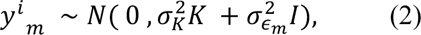

where 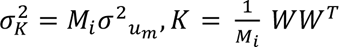. We will refer to *K* as the environmental kinship matrix, which represents the similarity, in terms of environmental covariates, between every pair of samples across grouping *i* (i.e., represent the correlation structure to the data).

#### Environmental explained variance (EEV)

The explained variance of a specific feature *y^i^_m_* environmental measurements by the

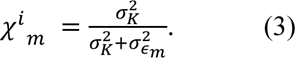

#### Estimating effective population size

To approximate effective population size (N_e_) we use fourfold degenerate synonymous diversity. Calculation was adapted from Schloissnig et al.(*44*):

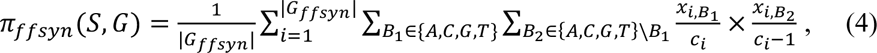

where S is the sample, G is the gene of interest, |G_ffsyn_| is the number of fourfold degenerate sites, *x_i_*,*B_i_* is the number of times nucleotide *B_j_* was seen in the fourfold degenerate position *i*, and *c_i_* is the coverage of position *i*.

#### Controlling for the effects of time and effective population size

To control for the effects of time and effective population size, we added these two potential confounders to our LMM as fixed effects. We formulate our LMM as follows:

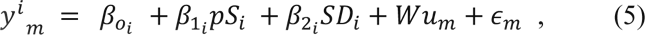

where *β*_0*i*_ is the intercept, pS (as a proxy for time) is the first fixed effect with *β*1*_i_* as its coefficient, *SD_i_* (as a proxy for effective population size) is the second fixed effect with *β*_2*i*_ as its coefficient. *W*is a normalized matrix of environmental factors, *u_m_* and *ϵ_m_* are independent random variables distributed as 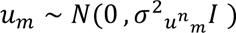 and 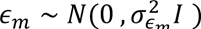. The parameters of the model are *β_oi_*, *β*1*_i_*, *β*2*_i_ σ*^2^*u_m_* and 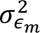

#### LMM on samples from the epipelagic zone

To demonstrate that the association of pN/pS with the environment is not confounded by depth, we reran the LMM (eq. 5) only on samples from the epipelagic zone (up to 76m, median depth across 746 samples).

#### Linear model for dN/dS rates

We modeled the benchmarking data by constructing a linear model for dN/dS (the dependent variable) with the environmental factors as independent variables:

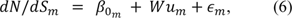

where *β*_0*m*_ is the intercept, *W* is a normalized matrix of environmental factors and *u_m_* is a vector of environmental coefficients. *ϵ_m_* ∼ *N*(0, *σ* ^2^*ϵ_m_ I*). *m* ∈ {1, …, *M*}, where M is the number of genes tested.

To account for the effect of time as potential confounder we constructed the following linear model for dN/dS (the dependent variable) with the environmental factors and dS (as a proxy for time) as independent variables.

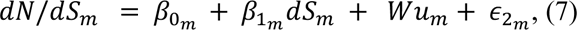

Where *β*_0*m*_ is the intercept, dS(as a proxy for time) is fixed effect with *β*_1*m*_ as its coefficient, *W* is a normalized matrix of environmental factors and *u_m_* is a vector of environmental coefficients that were available with these samples (phosphate, depth, nitrate + nitrite). *ϵ*_2*m*_ ∼ *N*(0, *σ*^2^ *ϵ*_2*m*_ *I*). *m* ∈{1, …, *M*}, where M is the number of orthologs tested. In this model we only used depth, NO_2_+NO_3_ and phosphate, as these were the environmental factors available for the benchmarking data.

To account for potential underlying structure in terms of phylogeny, we applied phylogenetic generalized least squares (*45*) (PGLS) regression in addition to ordinary least squares (OLS) regression to model dN/dS. Phylogenetic regression models are used when there is an underlying phylogenetic clustering structure of the observations. In such case, samples may be lowly or highly phylogenetically diverged leading to residuals that are not *i.i.d.*, which can affect the fit of the model. On the other hand, the caveat of PGLS is an underestimation of the *R*^2^ as compared to linear models (46, 47). In our data PGLS and OLS essentially gave the same results and we therefore only report the latter.

#### KEGG KO expression data as a ranking metric

Using expression data from 4,092 KEGG KOs collected by Kolody et al. (*22*), we ranked the KO genes in our marine samples in the following way. We first represent the expression data in relative abundance space (normalize each sample by its read counts). Next, for each focal KO *i*, where *i* ∈ {1, …, *I*}, we sum across its different instances within each sample. The input is an *m* × *n* matrix, where *m* is the number of different instances of KO *i*, and *n* is the number of samples. The output is a vector of length *n*: (*x*_1_, …, *x_n_*). Finally, we average the expression levels (*x*_1_, …, *x_n_*) across samples: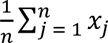 and rank the KOs based on the calculated average expression. We limited the scope of our analysis to samples collected only in small fraction filters (0.22 *μm*).

#### Determination of extracellular genes

To determine extracellular gene groups, we searched the eggNOG v.5 OG database for the words ‘secreted’ or both words ‘extracellular’ and ‘protein’ in their description. We demanded that the words ‘autoinducer’, ‘expression’, ‘role’ and ‘hypothetical’ are not in the description to prevent instances where (a) the OG in question describes a hypothetical protein; and (b) where the OG produces a secreted particle but is not secreted by itself, as is the case with autoinducer producing genes. To ensure robust pN/pS calculations, descriptions encompassing 10 OGs or more were assigned a group name, while descriptions encompassing less than 10 OGs were grouped together.

#### Calculation of mutation flux in divergent nitrate concentrations

To calculate the differential rate of nutrient incorporation between environments, we compare, for each pair of codons *a* and *b*, the ratio of mutations that result in *a* becoming *b* and the ratio of reverse mutations, that result in *b* becoming *a*. We term this quantity mutation flux. To prevent simplex-related confounding effects, mutation flux is estimated using the log-odds ratio between the two (e.g., *log* (*p*(*AAA* → *AAC*)/*p*(*AAC* → *AAA*)). We created a matrix *H*^(*U*)^ as follows: consider a set *U* of all genes for which SNP measurements exist and a subset *T* ⊆ *U* of this set across *K*samples. Let *G*^(*T*)^*j* = (*V*, *E*) be a codon graph for subset *T* and sample *j*, where *v* ∈ *V* is a codon (e.g., CUU coding for Alanine) and (*v*, *v*′) ∈ *E* if and only if *v* and *v*′ are one mutation apart (e.g., CUU for Alanine and CAU for Histidine). Let *w*^(*T*)^*j* : (*v*, *v*′) → [0,1] be a weight function where

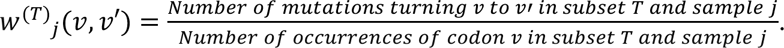

Let *H*^(*T*)^ be a matrix of dimension (|*E*| × *K*), where *H*^(T)^ (*v,v*’),*j* = *w*^(*T*)^ (*v*, *v*′). We next sum-normalized H per each sample and compared the codon mutation frequencies between the 40 lowest- and 40 highest-nitrate samples. Despite significant differences in codon mutation frequencies between low-nitrate and high-nitrate samples, some of the difference could be driven by the simplex properties of the sum-normalized codon mutation frequencies, and some could be attributed to the different rates of synonymous mutations between the high- and low-nitrate groups, which, combined with simplex properties, may affect observed nonsynonymous mutation rates. To address these simplex properties, we employed a centered log-ratio (CLR) normalization on *H*. The CLR transformation is a mapping, per codon composition, from the simplex to a Euclidean vector subspace. This log transforms each value and then centers them around zero as given below:

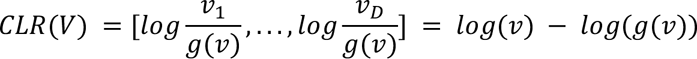

where g(v) is the geometric mean of all of the codons.

To address differences in rates of the different types of mutations, for each mutation (*v*, *v*′) in the CLR normalized matrix *H*′^(*U*)^ we calculated the log odds ratio between the mutation and its reverse mutation. Namely, we computed mutation flux matrix *F*^(*U*)^ where 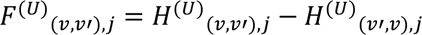. We compared differences in codon mutation flux between low- and high nitrate samples using the Mann-Whitney *U* test.

#### Calculation of expected random mutation cost per genetic code

Let *V*be a genetic code with a set *V*_s_⊂ *V*of stop codons. Let *P*(*v*) be the abundance of codon *v* ∈ *V* in a sample and *P*(*mut*(*v*, *v*′)) the probability of a single mutation from codon *v* to *v*′. Let *c_e_*: *V* × *V* →[0,1] be a cost function for element *e*, where:

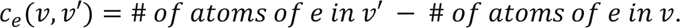

With these notations, we define the expected cost of genetic code *V* for element *e* as follows:

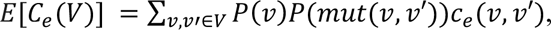

and the ERMC cost as:

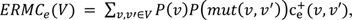

where *c*^+^*_e_* = *max*(0, *c_e_*). When testing the random mutation cost on hydrophobicity or PR we consider the absolute difference in the amino-acid hydropathy index or PR value pre- and post-mutation.

We estimate *ERMC_e_* (*V*) as follows:

1. We define *P*(*v*) as the median abundance of all codons *v*’ coding for the same amino acid as *v*.
2. We calculate mutation rates in sites that are under minimal selection. To this end, we estimate *P*(*mut*(*v*, *v*′)) by calculating, from fourfold-degenerate synonymous mutation sites the average abundance of each single nucleotide mutation (e.g. A to C) across all genes in which there are called SNPs in all ocean samples, excluding stop codons. We then estimate *P*(*mut*(*v*, *v*′)) using the relative abundances of all pairs of single nucleotide mutations. We estimate *P*(*mut*(*v*, *v*′)) for *Prochlorococcus* and *Synechococcus* using published transition:transversion rates (*1, 48*).
3. We calculate *c* using information on the amino acids which each codon codes for.

To compute a P-value, we generate a null distribution by calculating *ERMC_e_* (*V*) for hypothetical genetic codes. We randomize the first and second position of all codons, while maintaining that the two sets of first and second positions in which the stop codons reside are separated by a single transition mutation.

#### Confounding effects between cost functions for the structure of the genetic code

To confirm that our elemental cost function is not confounded by traditional properties of amino acids such as the polar requirement (PR) and hydropathy index (*11, 15*), we calculated the expected random mutation cost (ERMC), per hypothetical genetic code, using these common cost functions across 1 million codes. To this end, we randomized the first and second position of all codons, while maintaining that the two sets of first and second positions in which the stop codons reside are separated by a single transition mutation. We next calculated a contingency table for each pair of cost functions for both nitrogen and carbon (i.e., *ERMC_N_* (*V*):*ERMC_PR_* (*V*), *ERMC_N_* (*V*):*ERMC_hyd_* (*V*), (*V*):*ERMC_PR_* (*V*), *ERMC_C_* (*V*):*ERMC_hyd_* (*V*)). We assign each code to one of four bins in the following way: (1) surpassing the standard code in both cost functions (e.g., nitrogen and PR), (2) surpassing the standard code only in element *e* cost (e.g. only nitrogen), (3) surpassing the standard genetic only in the traditional cost function (e.g., PR), (4) not surpassing the standard code in either. We applied the Chi-square test of independence with two degrees of freedom to each contingency table.

Finally, we examined the subset of genetic codes (out of the 1M hypothetical codes) that have a lower ERMC than the standard code in terms of PR (*ERMC_PR_*) or hydropathy (*ERMC_hyd_*), and tested whether this subset is also optimized for nitrogen and carbon. To this end, we created a null model in which we compute EMRC only accounting for hydropathy or PR robustness and an expanded, nested model that adds minimization of nitrogen and carbon utilization. Under the null, we expect that in the subset of codes optimizing for hydropathy and PR, nitrogen and carbon utilization will also be minimized. Under the alternative, if such nutrient optimization indeed exists separately from structural optimization, we expect that nitrogen carbon and utilization will not be optimized. Exact P-values are defined to be the fraction of shuffled codes with ERMC smaller or equal to the standard code (one for each element - carbon and nitrogen) in this subset of codes that have a lower ERMC than the standard code in terms of PR (*ERMC_PR_*) or hydropathy (*ERMC_hyd_*).

#### Determination of ‘square’ and ‘diagonal’ arrangements of nitrogen in the genetic code

We define a ‘square’ arrangement of the codons coding for nitrogen-rich amino acids histidine, glutamine, asparagine, lysine and arginine as one where their codons span only two nucleotides in the first position and two nucleotides in the second position. In the standard code, these amino acids are coded by CAN, CGN, AAN, AGR, following a square configuration. In contrast, a ‘diagonal’ arrangement of the codons coding for these amino acids is one where they span all possible nucleotides in the first position and all possible nucleotides in the second position. For example, a genetic code where TTY codes for histidine, TTR for glutamine, CCN and AAR for arginine, GGY for asparagine and GGR for lysine constitutes a ‘diagonal’ arrangement of nitrogen amino acids. In each of the alternative genetic codes we generated, we tested whether either of these conditions hold and, if so, designated the code as ‘square’ or ‘diagonal’ accordingly.

#### *Prochlorococcus* and *Synechococcus* genomic data and mutation rates

We downloaded *Prochlorococcus* and *Synechococcus* protein-coding gene sequences (where available) from the Joint Genome Institute (https://genome.jgi.doe.gov/portal/) following accession numbers published by Berube et al. (1). To estimate codon relative abundance *P*(*v*), we counted and sum-normalized codons in all protein-coding genes for each strain. To estimate codon mutation rate *P*(*mut*(*v*, *v*′)), we used the published transition:transversion rate of 2:1 for Prochlorococcus and *Synechococcus* (*23*).

#### Multiple taxa ERMC calculation

To calculate ERMC for 39 taxa across multiple transition:transversion rates, we downloaded codon usage and GC-content data collected by Athey et al. (*49*). We used codon usage counts to estimate *P*(*v*) and 11 transition:transversion rates (1:5, 1:4, 1:3, 1:2, 2:3, 1:1, 3:2, 2:1, 3:1, 4:1, 5:1) to estimate *P*(*mut*(*v*, *v*′)).

### Supplementary Text

#### Supplementary Note 1. Conservative nonsynonymous substitutions are more common than radical ones in marine microbial metagenomes

To corroborate the validity of calculating selection metrics from metagenomic samples, we sought to replicate a known property of mutations in amino acids, namely that mutations resulting in “conservative” amino acid replacements are more common than those resulting in “radical” ones (*50*). To this end, we calculated the ratio of nonsynonymous mutations leading to radical amino acid replacements and compared them to those leading to conservative replacements. To best define “radical” and “conservative” nonsynonymous substitutions, we used two known amino acid properties: (1) the Polar Requirement (PR) scale and (2) amino-acid hydropathy index (**Table S2**). Since the definition of radical and conservative may be subjective, we considered multiple thresholds as radical and conservative for each property. For example, a radical substitution could be one that results in a difference of more than 3 units on the hydropathy index whereas a “conservative” one results in just one or less (**Table S2**; hydropathy case 4).

We then calculated, for each KEGG ortholog and for each radical/conservative threshold, the proportion of radical nonsynonymous substitutions (out of all possible radical sites) as compared to the proportion of conservative ones across samples using a Mann-Whitney U test. P-values were adjusted to control for multiple hypotheses.

Formally, our statistical analysis postulates:

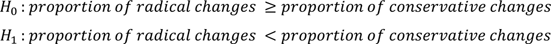

Next we calculated, for each radical/conservative threshold, the proportion of KEGG orthologs in which we reject the null hypothesis (i.e., the proportion of KOs in which nonsynonymous mutations leading to conservative amino acid replacements are significantly more common than nonsynonymous mutations leading to radical amino acid replacements) and conducted a permutation test. We permuted the sample labels of each radical/conservative threshold across 10,000 iterations, where in each iteration we calculated the proportion of KEGG orthologs in which we reject the null hypothesis. The P-value, per radical/conservative threshold, is defined to be the proportion of iterations in which the ratio of KEGG orthologs in which we reject the null hypothesis is higher as compared to the original data. We find that these comparisons are significant for all cases, regardless of the amino acid property in question or the implemented threshold and that this is the case for most KOs (P<0.0001 for all comparisons; **Table S2**). Finally, we examined whether highly expressed genes have fewer radical amino acid replacements as compared to lowly expressed genes. To this end, we used an additional expression dataset for marine microbial gene expression to rank KEGG orthologs by their mean expression (similar to the approach implemented in **Fig. 1D** (*22*)). We find that KEGG orthologs which exhibit significantly more conservative mutations than radical ones have significantly higher expression as compared to the alternative case (Wilcoxon signed-rank test P<10^-5^). These results demonstrate that metagenomic sequences are adequate for analyzing selective pressures.

#### Supplementary Note 2. Resource-driven selection drives specific amino acid changes

Our results indicate that mutations leading to an addition of resources would be more common in nutrient-rich environments. This is supported by previous observations regarding genomic and proteomic changes associated with environmental concentrations of nitrate (*2, 5, 21, 51–53*). We thus sought to quantify how resource conservation manifests in molecular changes to DNA and protein sequences. Mutations in nitrogen-rich and, typically, carbon-poor environments were shown to drive an increase in genomic GC content (which increases nitrogen requirements), as well as by higher nitrogen and lower carbon in protein sequences.

Using SNP derived from metagenomic data, we corroborate previous findings (*5, 21*), and show that mutations leading to higher GC content and those leading to higher nitrogen proteins are more prevalent in high-nitrate waters as compared to low-nitrate (**Fig. S12A,B**). We note that mutations leading to higher GC content (higher nitrogen DNA) often result in higher nitrogen proteins (*2, 3*). We further deconvolve these two types of mutations and find that the ratio of mutations that lead to higher nitrogen DNA but do not change protein nitrogen-content increases with environmental nitrate (R=0.333; **Fig. S12C**). Additionally, the ratio of mutations that increase protein nitrogen but do not change DNA nitrogen-content is even more strongly correlated with nitrate concentrations (R=0.473; **Fig. S12D**). We hypothesize that this stronger association of the environment with protein sequences can be attributed to the higher copy number of protein as compared to DNA and RNA, often in the order of 10^2^-10^5^ protein copies per RNA molecule (*54, 55*) leading to higher cost per gene, and thus to stronger selective pressure. Our results suggest a differential effect of mutations on nutrient incorporation in DNA as compared to protein sequences.

To study amino acid substitutions typical to low- and high-nitrate environments, we examined mutations leading to changes in protein nutrient content by comparing codon mutation frequencies in low- and high-nitrate samples, after accounting for simplex-related confounders (Methods). We found significant differences in codon mutation frequencies between environments (**Fig. S13A-C**). We next sought to examine the typical change in nutrient consumption in varying nitrate concentrations. We therefore defined mutation flux as the ratio between a codon mutation and its reverse, thereby measuring the typical flux of nutrients; and estimated it using the log-odds ratio between the two (e.g., *log* (*p*(*AAA* → *AAC*)/*p*(*AAC* → *AAA*)). Notably, across all the mutations significantly more prevalent in samples from high nitrate environments (Methods), averaged across amino acids, we find a total increase in nitrogen (**Fig. S13D**; 18 N atoms summed across all significant amino acid changes, P=0.0082), decrease in carbon (**Fig S13D**; -37 atoms, P=0.0165), decrease in sulfur (-6 atoms, P=0.009), a decrease in molecular weight (-508.91 g/mol, P=0.0193) and a non-significant decrease in oxygen (-6 atoms, P=0.1505). These results provide additional evidence that the higher the nitrate concentrations are, the weaker the selection against mutations leading to higher nitrogen incorporation into protein sequences. While nitrate concentrations increase with depth (**Fig. S2I**), labile dissolved organic carbon concentrations typically decrease (*56*). We therefore also show the specific codon mutations reducing carbon incorporation into proteins in nitrogen rich and typically carbon poor, environments (**Fig. S13D,E**).

#### Supplementary Note 3. The association between the environment and pN/pS is robust to time, effective population size and environmental niche

The magnitude of purifying selection, approximated by pN/pS ratios, may increase as a function of both time and effective population size (*57, 58*). These two mechanisms might also be associated with environmental conditions and we thus sought to control for them as potential confounders. Time may affect the magnitude of pN/pS ratios via random genetic drift. Effective population size may affect selection via competition, as in large populations, any slightly deleterious mutation is rapidly selected against. We used the ratio of synonymous substitutions (i.e., pS) calculated from our data as an approximation of dS and representing neutral polymorphism rate, as a proxy for time (*59*). To approximate effective population size, we relied on population genetic theory suggesting that effective population size is scaled to diversity at the silent sites when assuming constant mutation rate between populations. As our data is composed of closely related sequences and hence similar mutation rate, we used fourfold-degenerate synonymous diversity (SD) as a proxy for the effective population size (*44*) (Methods).

We first tested whether SD is associated with environmental conditions, and compared the SD of populations from different oceanic regions with different environmental parameters. We found that pN/pS was more strongly associated with most environmental parameters than SD (Kolmogorov-Smirnov test P<10^-20^ for depth, nitrate, oxygen, phosphate, silicate and temperature; P<10^-5^ for NO2+NO3 concentrations; non-significant for salinity; **Fig. S3**) suggesting that selection on protein sequences may be more strongly affected by the environment than effective population sizes are. We further found that in KOs with the highest EEV these differences are the most pronounced (Wilcoxon signed-rank test P<10^-15^; **Fig. S14**).

As LMMs also allow controlling for potential confounders as fixed effects, we next added our approximations of time and the effective population size to our LMM in order to account for their effect on the magnitude of pN/pS ratios. Accounting for these potential confounders as fixed effects would enable a more accurate quantification of the environmental effect. Similar to the results of our original LMM, across both KEGG and eggNOG orthologs, we found that a significant fraction of the variance in pN/pS can be attributed to the environment, even after controlling for these confounders (**Fig. S4C,D**; Mann-Whitney *U* test P<10^-16^). We found a high correlation between EEV estimated by the two models (R^2^=0.947 and 0.929 for KEGG and eggNOG, respectively; **Fig. S15**).

To demonstrate that our results are not confounded by depth, the environmental variable with the second strongest correlation to pN/pS (after nitrate), we recalculated our LMM only on samples from the epipelagic zone (Methods). Similar to the original results of the LMM, we show significant variance explained in pN/pS in this environmental niche (**Fig. S6**; Mann-Whitney *U* test P<10^-20^). Taken together, these results demonstrate that the association between the environment and pN/pS is robust to time, effective population size and environmental niche.

#### Supplementary Note 4. The association between environmental measurements and the magnitude of purifying selection is replicated using single cell data and dN/dS

pN/pS, the ratio of nonsynonymous polymorphisms (pN) to synonymous polymorphisms (pS), approximates dN/dS which quantifies the magnitude and type of selection (purifying, neutral or positive) exerted on protein-coding sequences across a phylogenetic group (*44, 60*). While dN/dS calculation requires genomic sequences at the strain-level that are available for only a small number of samples and from limited strains, it allows for a more accurate quantification of purifying selection in the taxa represented in the phylogeny. We therefore sought to confirm our findings using dN/dS. To this end, we used benchmarking data in the form of assembled genomes from uncultivated single cells from dominant and representative lineages of the surface ocean (SAR-11, SAR-86 and Prochlorococcus) (*20*), and their corresponding environmental measurements (Methods).

We calculated dN/dS for KEGG orthologs in each lineage. Given the small number of samples, we estimated the fraction of variance explained by the environment using a linear model with environmental covariates (Methods). Across all bacterioplankton lineages comprising the benchmarking data, we found that a significant portion of the variance in dN/dS ratios calculated from single-cell data is explained by the environment (SAR-11, SAR-86, Prochlorococcus; Wilcoxon signed-rank test P<10^-16^; **Fig. S4E-G**). Similar to our original LMM, we note that this association holds, across all three bacterioplankton lineages, even after controlling for both time as a confounder (Wilcoxon signed-rank test P<10^-16^; **Fig. S16**) and for phylogeny, using phylogenetic generalized least-square regression (Methods). These results validate our findings using pN/pS calculated from metagenomic sequences and demonstrate the association between selective pressure and environmental conditions, even in a single clade. This indicates that associations of pN/pS or dN/dS with environmental variables are likely not confounded by organismal properties which may differ in their metabolism across environments, but rather exhibit a consistent trend even within a single clade.

---

**Fig. S1.**
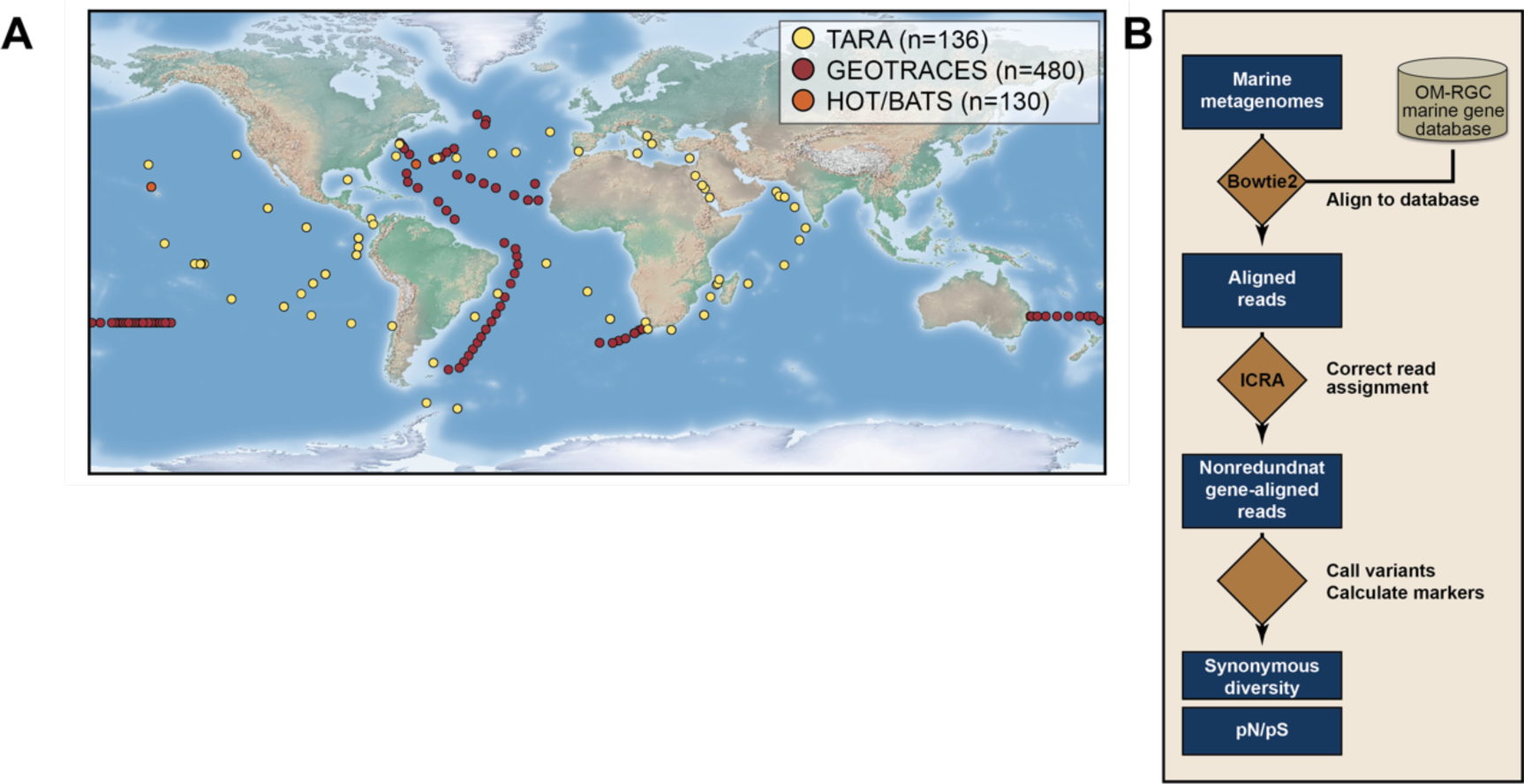
(A) Geographical overview of the samples used in this study. (B) Illustration of our computational pipeline.

**Fig. S2.**
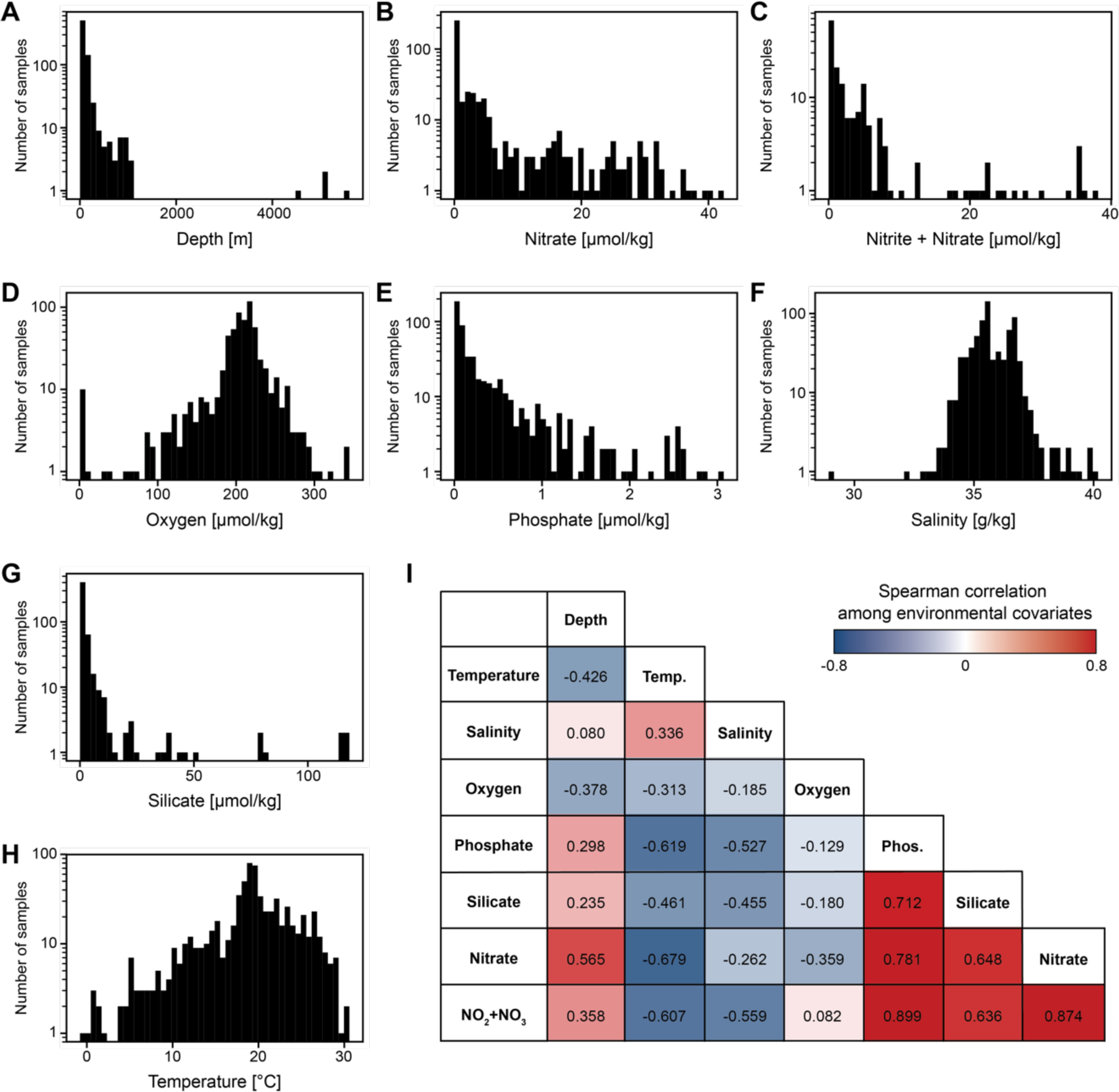
(A-H) Distribution of measurements taken alongside marine microbial samples for depth (A), nitrate (B), nitrate and nitrite (C), oxygen (D), phosphate (E), salinity (F), silicate (G) and temperature (H). (I) Spearman correlation coefficients between all pairs of environmental measurements across all available samples.

**Fig. S3.**
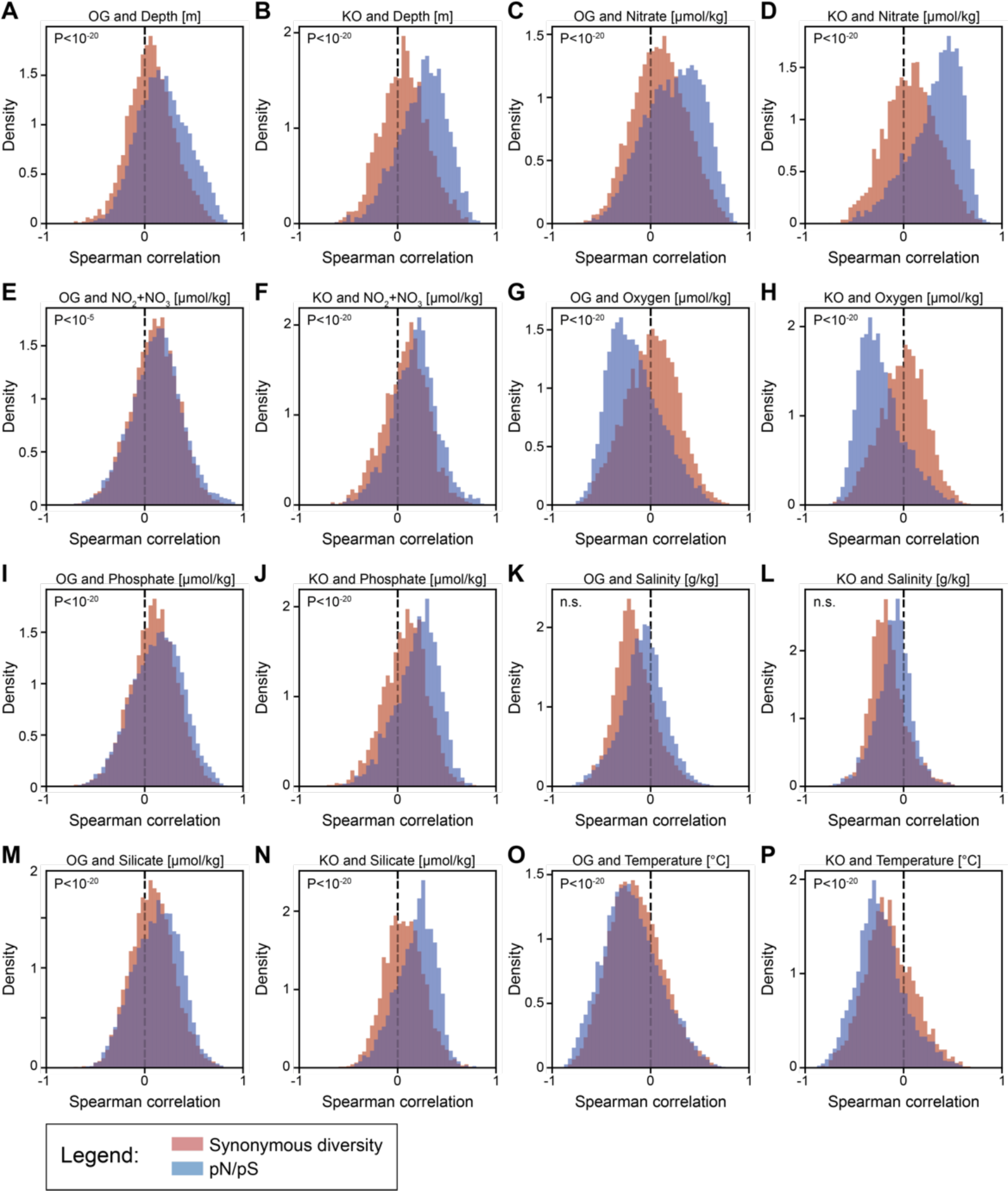
(A-P) Histograms of Spearman correlations between synonymous diversity (red) or pN/pS (blue) and environmental variables, for both KEGG KOs and eggNOG OGs. Panels A, C, E, G, I, K, M and O depict correlations between OG calculated parameters and depth, nitrate, nitrite and nitrite, oxygen, phosphate, salinity, silicate and temperature, respectively. Panels B, D, F, H, J, L, N and P depict correlations between KO calculated parameters and depth, nitrate, nitrite and nitrite, oxygen, phosphate, salinity, silicate and temperature, respectively. P, one-sided Kolmogorov-Smirnov test.

**Fig. S4.**
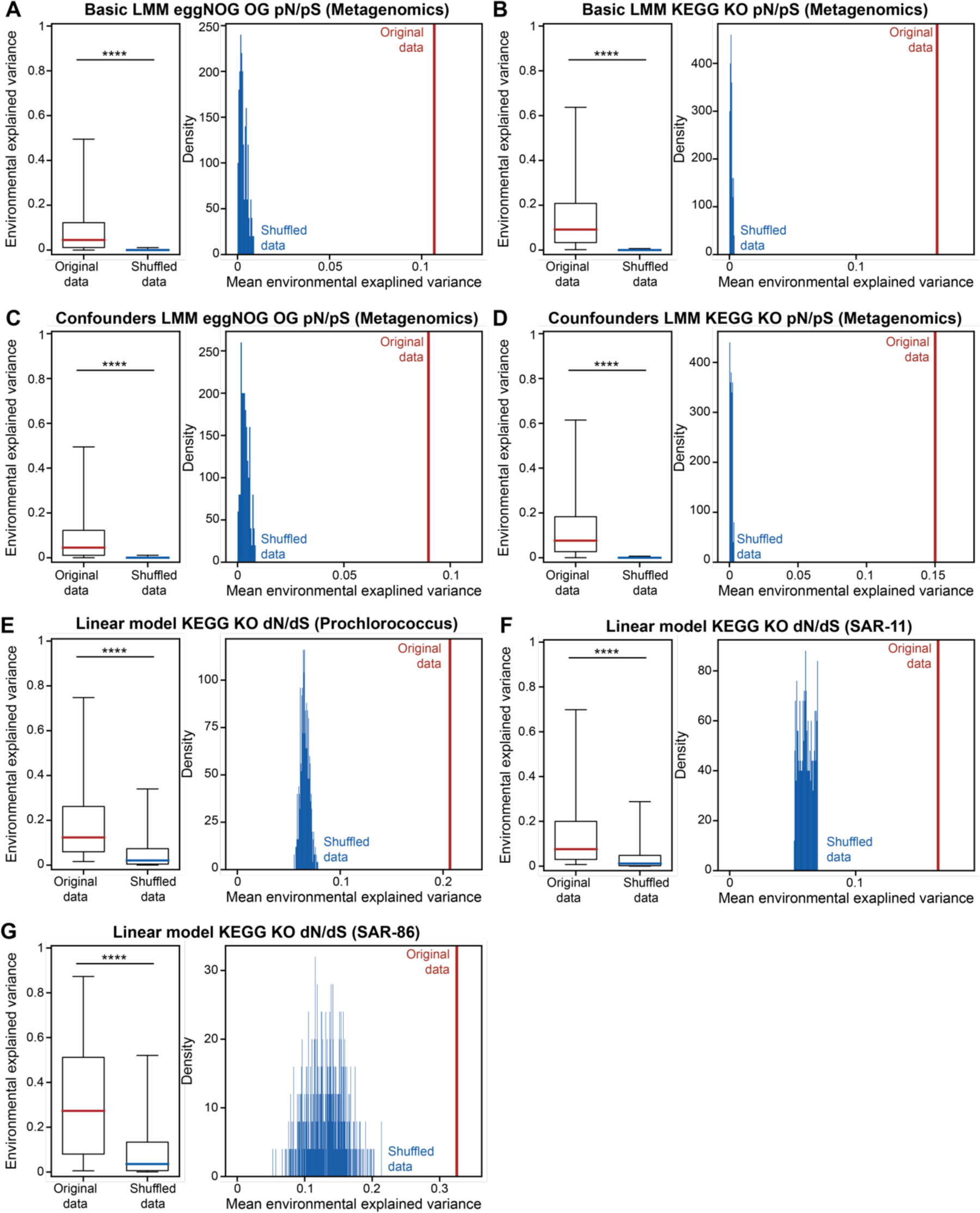
(A) Left box plot (line, median; box, IQR; whiskers, 5th and 95th percentiles) of the variance of eggNOG rates that was explained by the environment in a basic linear mixed model (LMM) setting as compared to the same data with shuffled labels; Right, mean variance explained in unshuffled data (red) as compared to a histogram (blue) of mean variance explained in 500 executions with shuffled data. (B) Same as A, for KEGG KO pN/pS. (C) Same as A, for variance in eggNOG OG pN/pS in a LMM corrected for confounding factors of time and effective population size. (D) Same as C, for KEGG KO pN/pS. (E) Box plot (line, median; box, IQR; whiskers, 5th and 95th percentiles) of the variance of KEGG KO dN/dS in genus *Prochlorococcus* that was explained by the environment in a linear model. (F) Same as E, for the SAR-11 clade. (G) Same as E, for the SAR-86 clade. ****, Wilcoxon signed-rank test P<10^-16^.

**Fig. S5.**
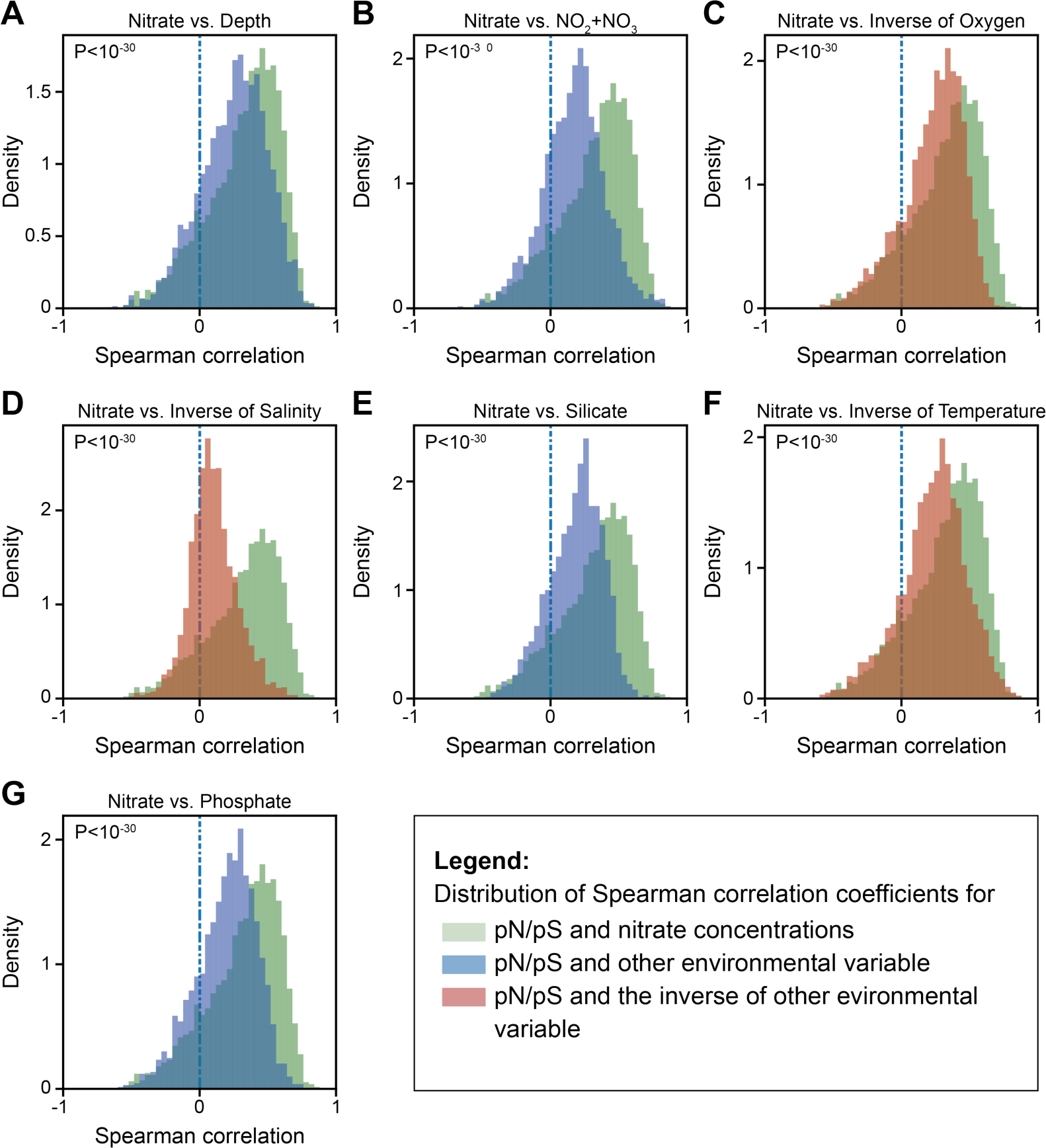
Histograms of Spearman correlations between environmental variables (blue) or their inverses (red) and pN/pS ratios of KEGG KOs as compared to a Spearman correlation between the concentration of nitrate and pN/pS ratios (green). P, one-sided Kolmogorov-Smirnov test. We note that unmeasured variables correlated with nitrate, such as labile dissolved organic carbon, may also be correlated with pN/pS.

**Fig. S6.**
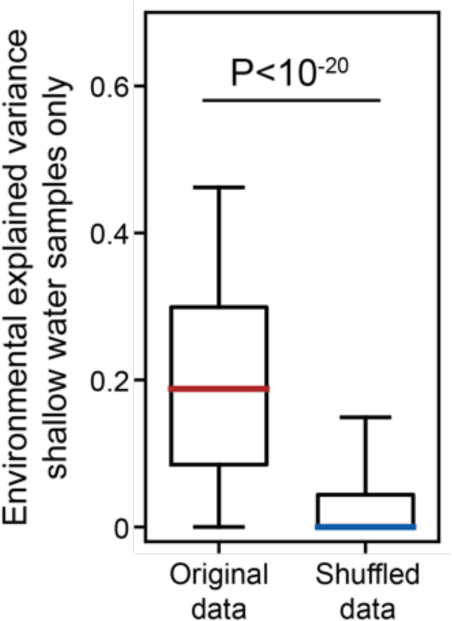
Box plot (line, median; box, IQR; whiskers, 5th and 95th percentiles) of the variance of KEGG KO pN/pS ratios that was explained by the environment in epipelagic samples (up to 76m) as compared to the same data with shuffled labels.

**Fig. S7.**
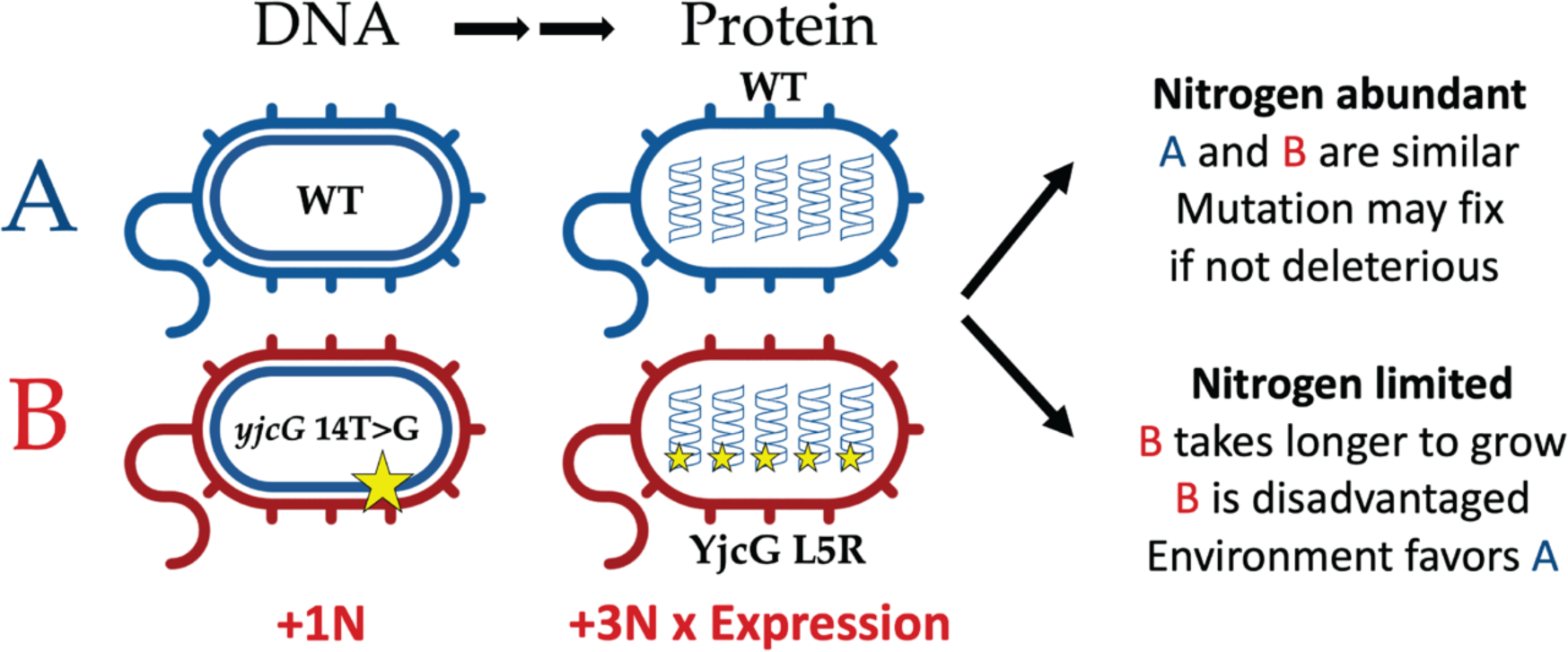
Illustration of the potential environmental effect of nitrate on protein-coding sequences. Depicted are two microbes, wild-type (WT; A) and a mutant with a T-to-G transversion in the 14th position of the yjcG gene, leading to a leucine to arginine substitution, thereby consuming three extra nitrogen atoms per protein molecule (B). If environmental nitrate is scarce and growth-limiting, A is likely to have a selective advantage over B, which over multiple generations would translate into an extinction of the B variant. In highly expressed genes such a mutation is likely to have an even stronger effect, leading to a quicker extinction. On the molecular level, this phenomenon would translate to lower pN/pS in nitrate-limited waters as compared to nitrate-rich ones.

**Fig. S8.**
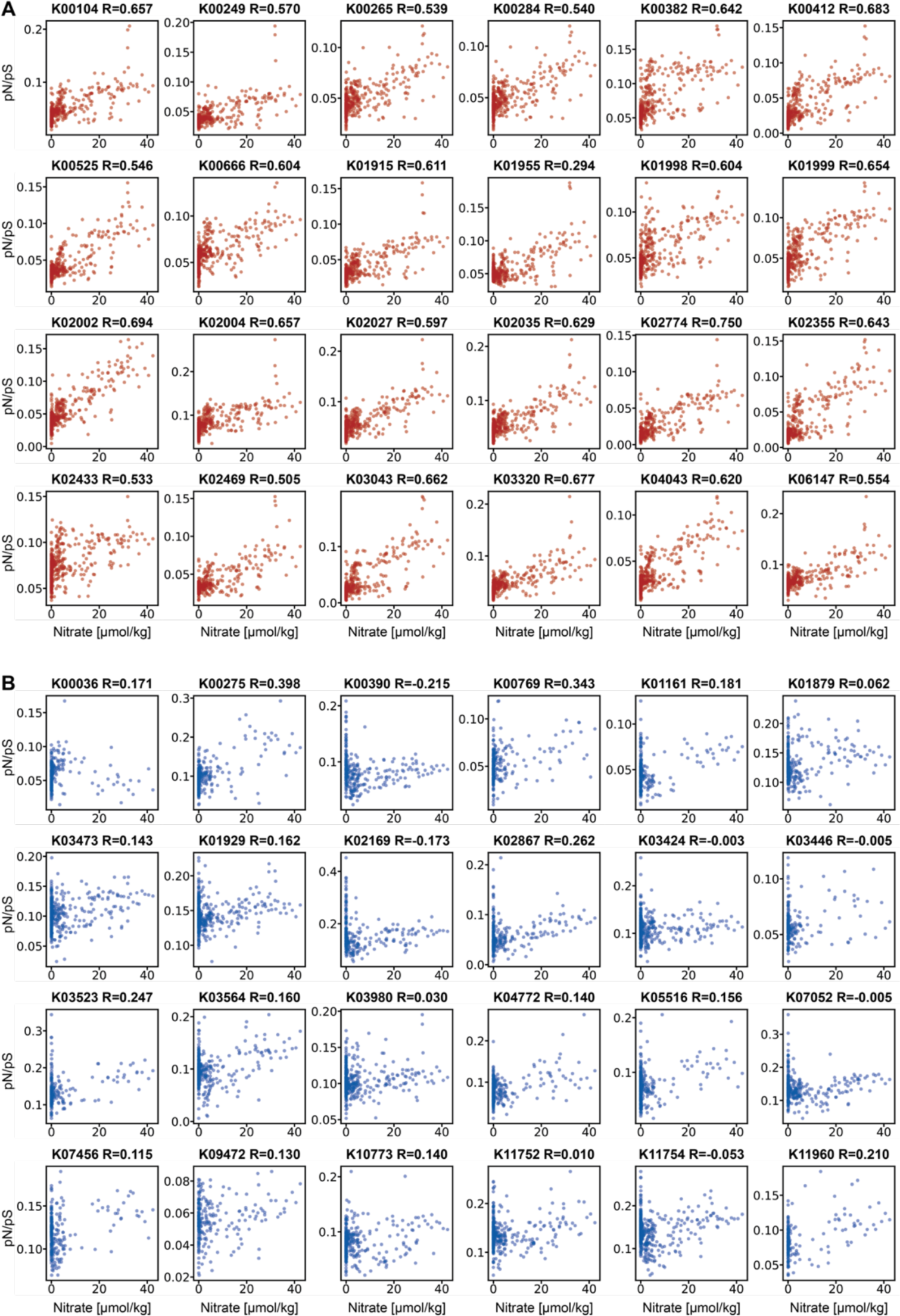
(top, red) 24 highly expressed KOs, exhibiting high correlation between pN/pS and nitrate concentrations. (bottom, blue) 24 lowly expressed KOs, exhibiting low correlation between pN/pS and nitrate concentrations. R, Spearman rho.

**Fig. S9.**
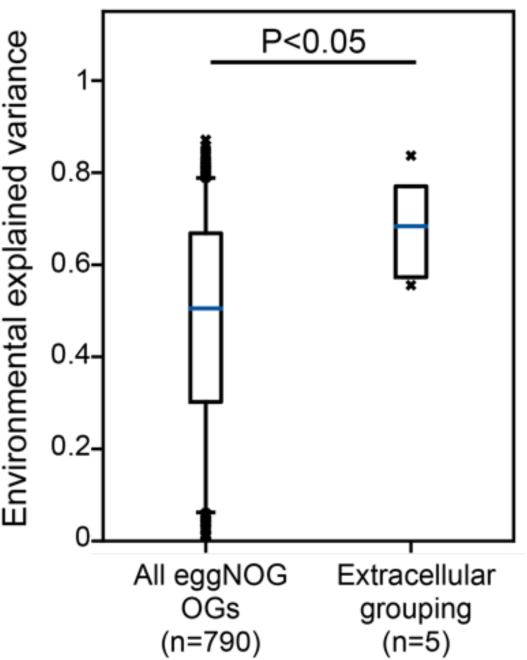
Box plot (line, median; box, IQR; whiskers, 5th and 95th percentiles) of variance in pN/pS explained by the environment in extracellular gene groups versus all eggNOG OGs.

**Fig. S10.**
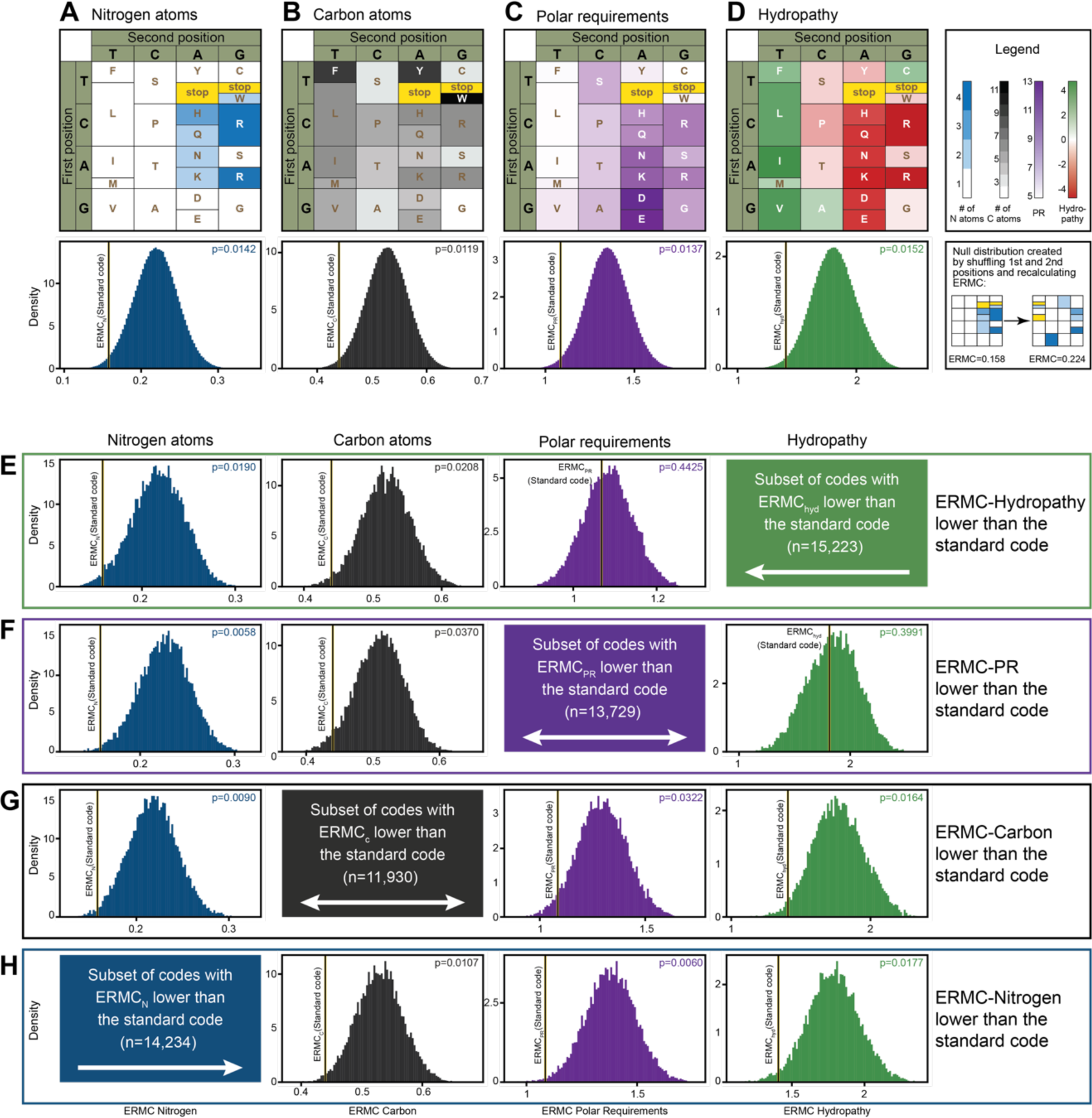
Upper panel: (Top) Nitrogen content (A; blue color scale), carbon content (B; black), the polar requirement (C; purple) and hydropathy (D; green-red) of different amino acids depicted along their positions in the standard genetic code. (Bottom) Histograms of the expected random mutation cost (ERMC), in 1,000,000 random permutations of the genetic code for nitrogen (blue), carbon (black), hydropathy (green) and polar requirements (purple). Lower panel: Histograms of ERMC for nitrogen (blue), carbon (black), hydropathy (green) and polar requirements (purple) for the subset of shuffled genetic codes with ERMC lower than the standard genetic code for (E) hydropathy, (F) the polar requirement, (G) carbon and (H) nitrogen. Vertical black-yellow line, ERMC of the standard genetic code for each property. P, permutation test.

**Fig. S11.**
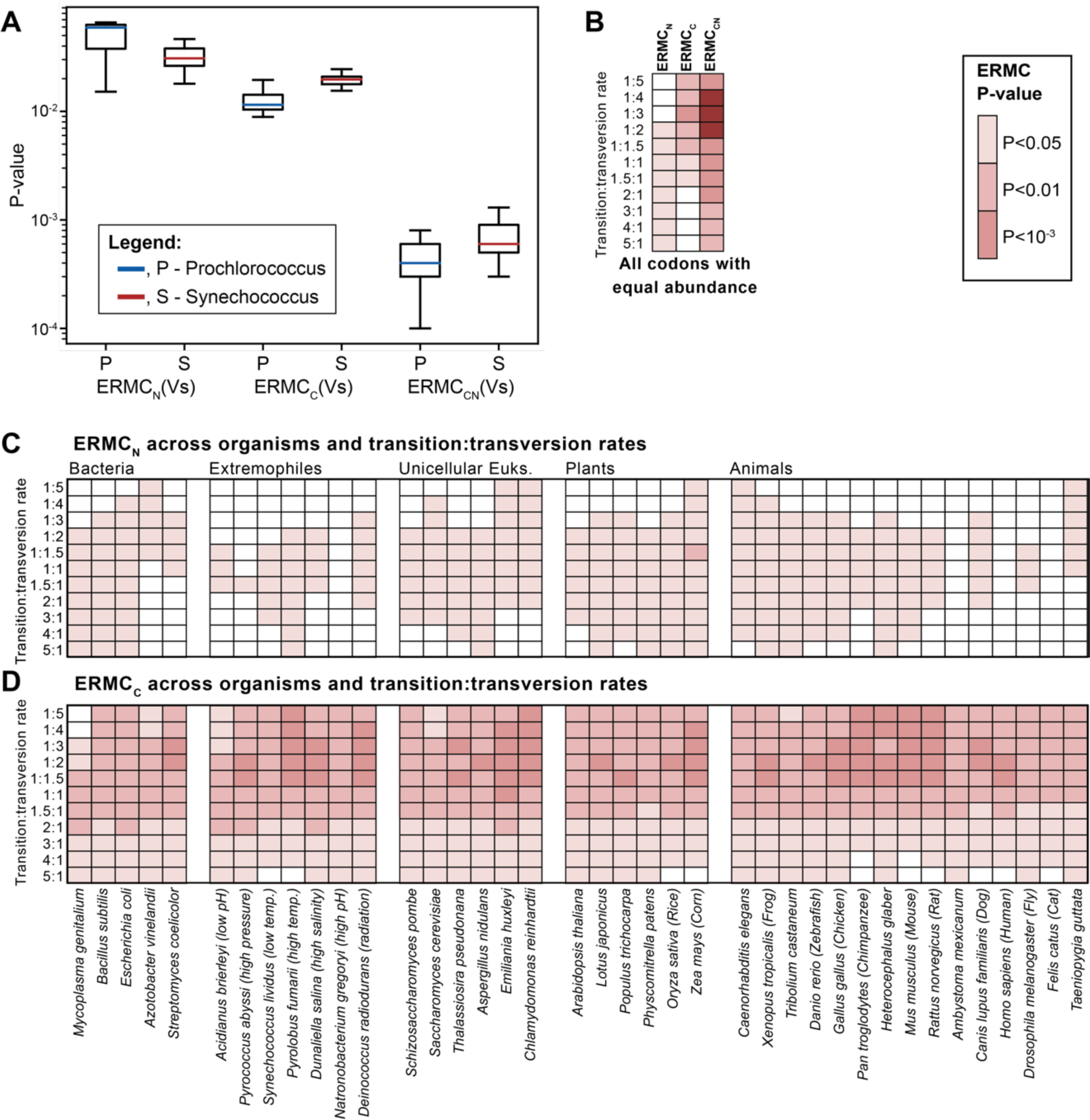
(A) Box plots (line, median; box, IQR; whiskers, 5th and 95th percentiles) of P-values for the ERMC of the standard genetic code for nitrogen (left), carbon (center) and both (right) across 187 Prochlorococcus (P, blue) and Synechococcus (S, red) strains. (B) Heat map of ERMC P-values for nitrogen, carbon and both, for a theoretical case in which all codons are of the same abundance. (C,D) Same as Fig. 2C for ERMCN (D) and ERMCC (E) P-values.

**Fig. S12.**
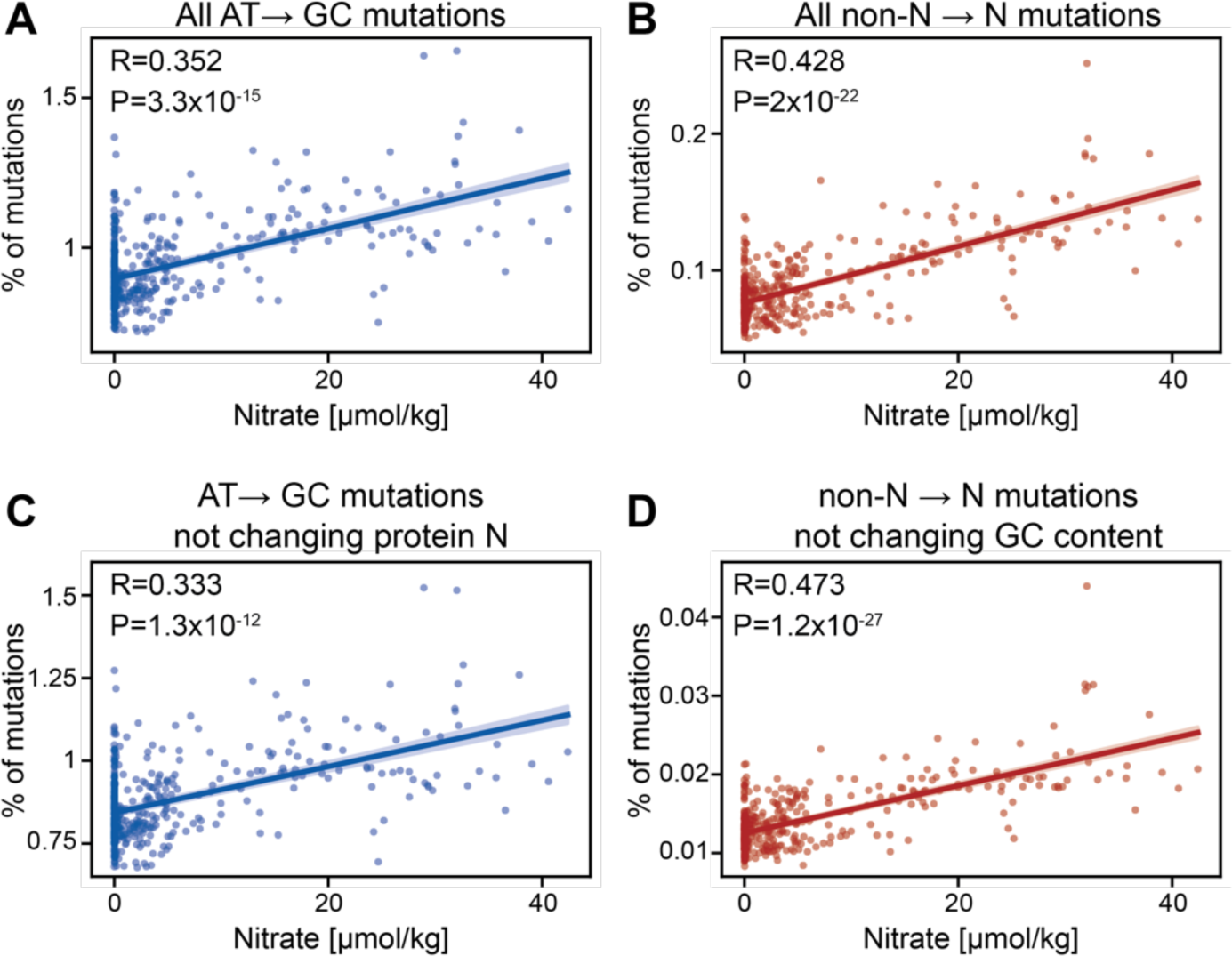
Scatter plots of nitrate concentrations (x-axis) versus percent of mutations (y-axis) for (A) all AT to GC mutations; (B) All mutations in coding sequences changing an amino acid with no nitrogen in its side-chain to one with nitrogen in the side-chain (non-N to N mutations); (C) AT to GC mutations not changing protein nitrogen; (D) non-N to N mutations not changing GC content. R and P, Spearman correlation coefficient and P-value; line, linear fit; shaded area, 95% confidence interval.

**Fig. S13.**
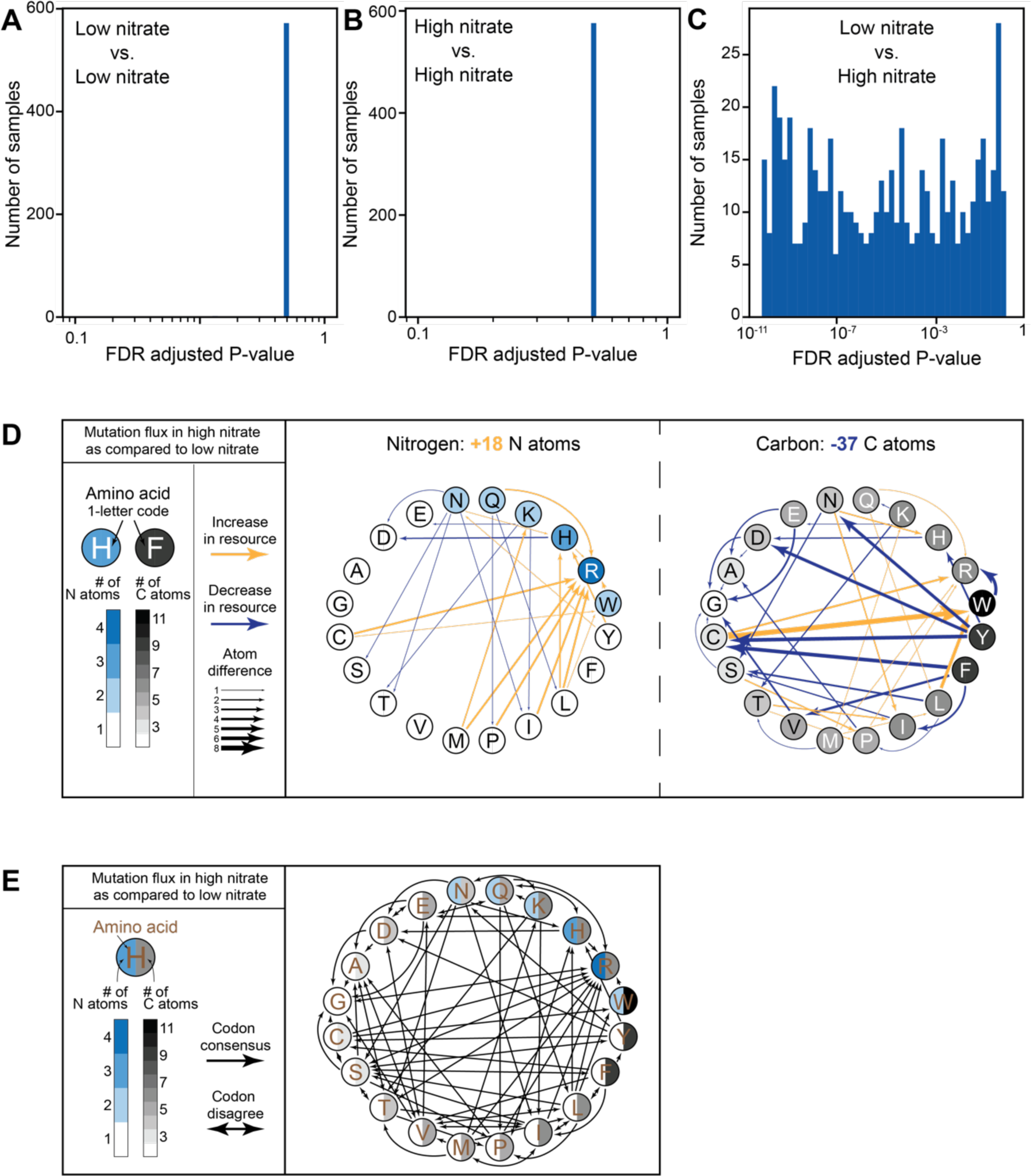
(A-C) Histograms of the distribution of Mann-Whitney *U* P-values of codon-to-codon mutations compared between (A) 40 low-nitrate samples and 40 other low-nitrate samples selected randomly out of the 80 lowest-nitrate samples; (B) 40 high-nitrate samples and 40 other high-nitrate samples selected randomly out of the 80 highest-nitrate samples; (C) 40 low-nitrate samples and 40 high-nitrate samples selected randomly out of the 80 lowest and highest nitrate samples. (D) Depiction of mutation flux (Methods) common in high versus low environmental nitrate concentrations, affecting amino acid left) and carbon (right) content. Yellow arrows, increase in resource; blue arrows, decrease in rrow thickness corresponds to number of atoms changed by mutation. (E) Same as D, showing regardless of change in resources. Two-headed arrows mark cases where different codons of mino acid have opposite mutation fluxes.

**Fig. S14.**
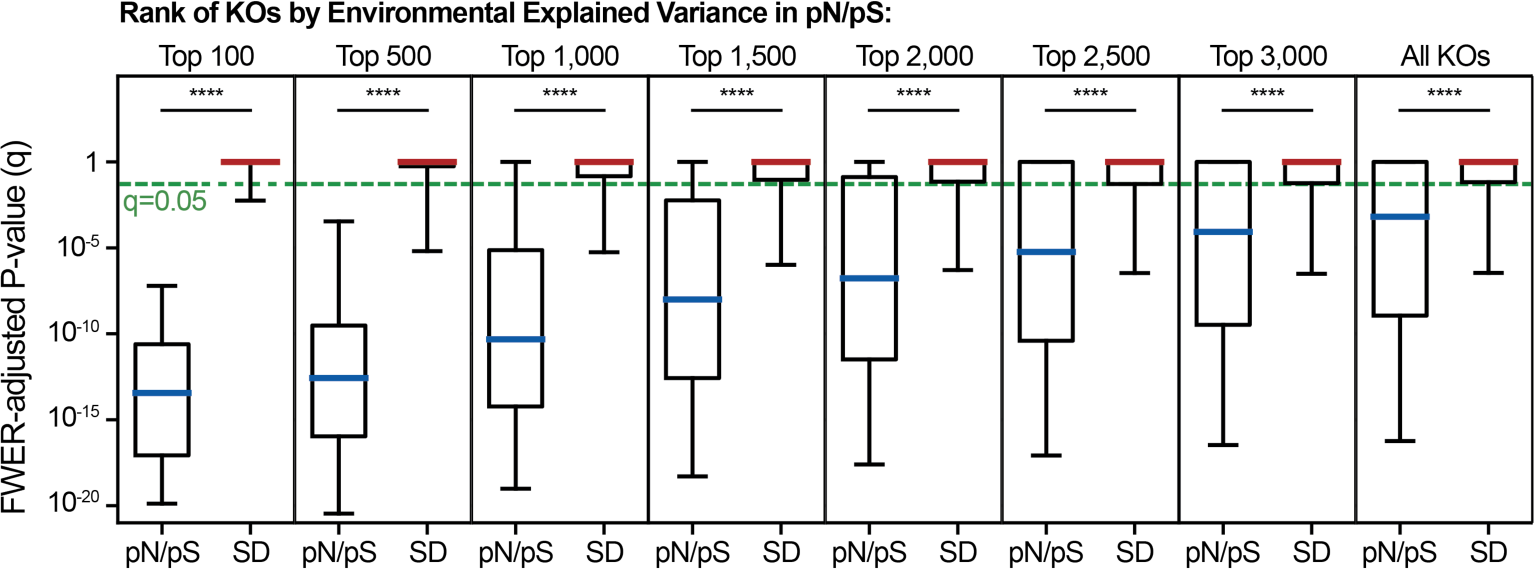
Box plots (line, median; box, IQR; whiskers, 10th and 90th percentiles) of P-values (t-test, multiple hypothesis adjustment) between samples with high-N (more than 19.6 µmol/kg) and low-N (0 µmol/kg), for KEGG KOs ordered by their EEV in the LMM, for both pN/pS (blue) and synonymous diversity (red). ****, Wilcoxon signed-rank test P<10^-15^.

**Fig. S15.**
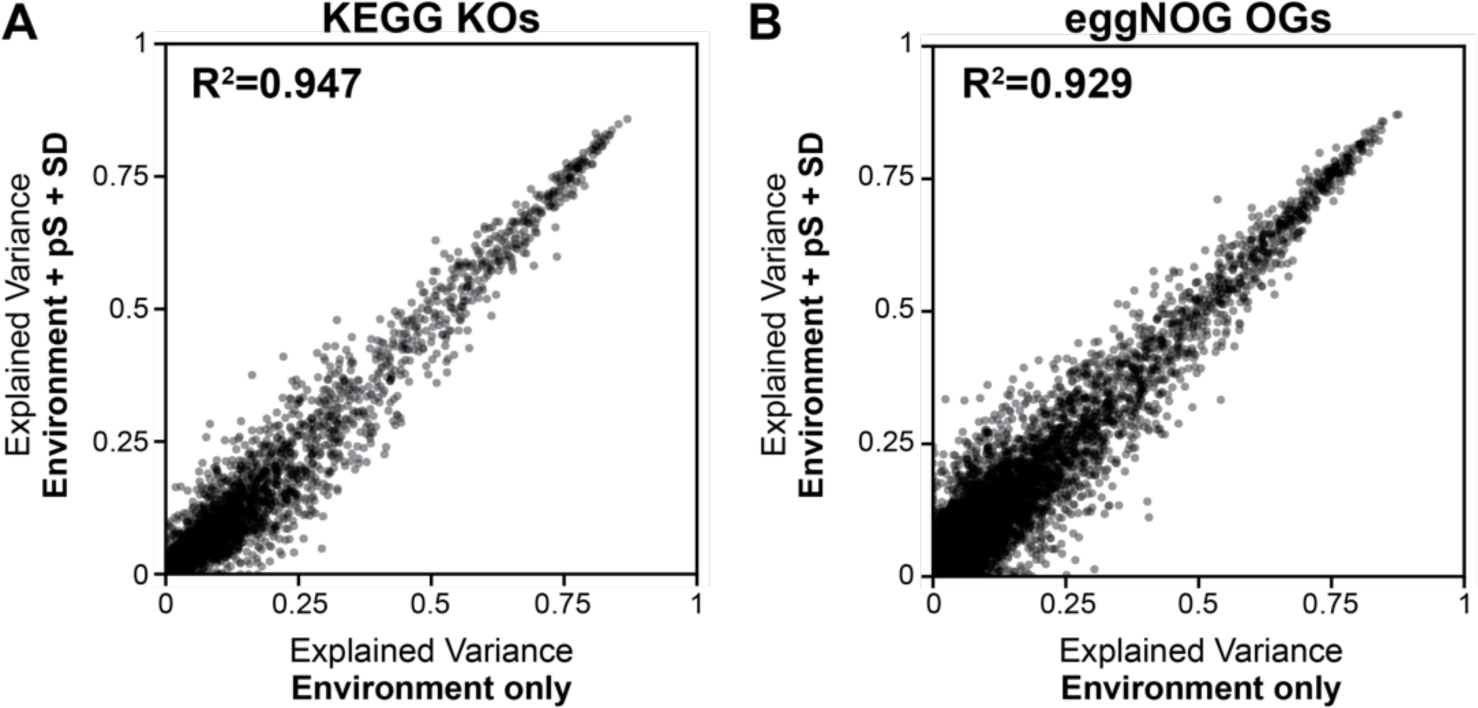
(A,B) scatter plot of explained variance in a linear mixed model considering only environmental factors as random effects (x-axis) versus a model considering environmental factors as random effects plus dS (as a proxy for time) and synonymous diversity (as a proxy for effective population size) as fixed effects (y-axis) in all (A) KEGG KOs and (B) eggNOG OGs.

**Fig. S16.**
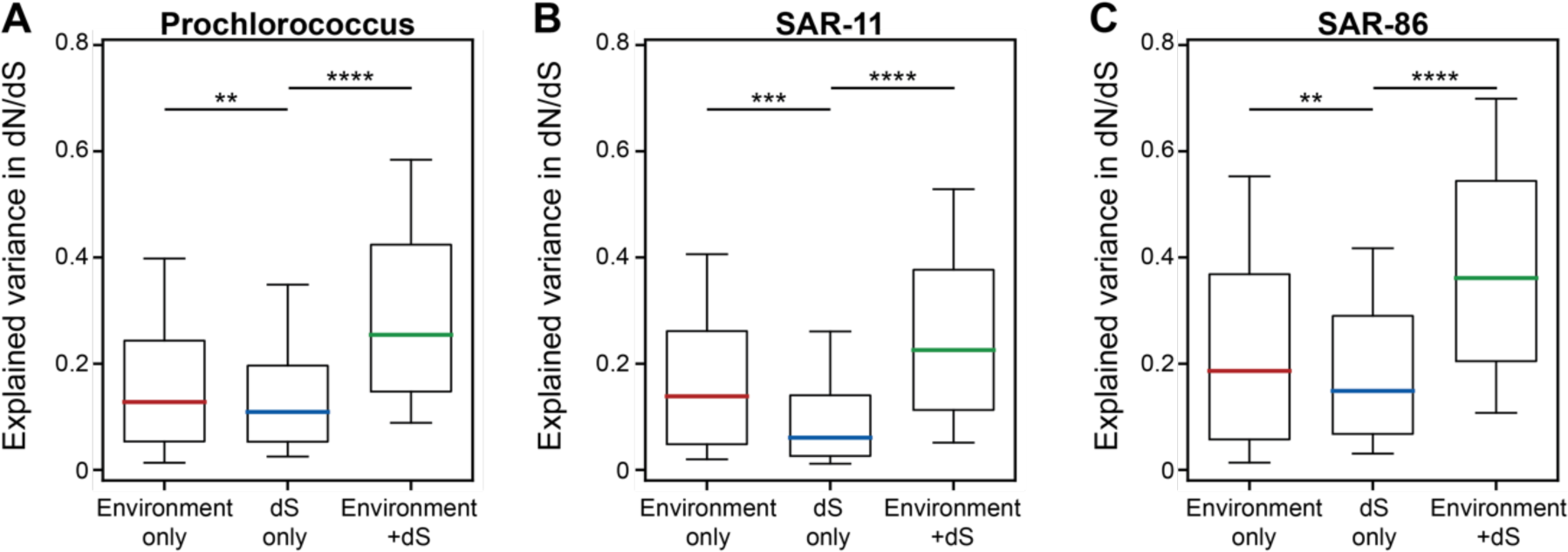
Box plots (line, median; box, IQR; whiskers, 10th and 90th percentiles) of explained variance in dN/dS using linear models considering the environment only (left, red), dS only (middle, blue), or both (right, green). (A) Prochlorococcus, (B) SAR-11 clade and (C) SAR-86 clade. **, Wilcoxon signed-rank test P<0.01; ***, P<10^-5^; ****, P<10^-16^.

**Table S1.** Contingency tables for each pair of cost functions for both nitrogen (n+) and carbon (c+), compared to PR (pr) and Hydropathy index (hyd), across 1 million simulated genetic codes. Each code is assigned to one of four bins: (1) surpassing the standard genetic code in both cost functions (<; <), (2) surpassing the standard genetic code only in element e cost (<; >=), (3) surpassing the standard genetic code only in the traditional cost function (>=; <), (4) not surpassing the standard genetic code in neither (>=; >=). Chi-square test of independence was applied to each contingency table.

| Cost function a | Cost function b | Sign a | Sign b | Alternative code counts | Chi-squared P-value |
| --- | --- | --- | --- | --- | --- |
| n+ | c+ | < | < | 128 | 1.32E-03 |
|  |  |  | >= | 14106 |  |
|  |  | >= | < | 11802 |  |
|  |  |  | >= | 973964 |  |
| n+ | hyd | < | < | 270 | 2.71E-04 |
|  |  |  | >= | 13964 |  |
|  |  | >= | < | 14953 |  |
|  |  |  | >= | 970813 |  |
| n+ | pr | < | < | 83 | 4.68E-16 |
|  |  |  | >= | 14151 |  |
|  |  | >= | < | 13646 |  |
|  |  |  | >= | 972120 |  |
| c+ | hyd | < | < | 249 | 4.86E-07 |
|  |  |  | >= | 11681 |  |
|  |  | >= | < | 14974 |  |
|  |  |  | >= | 973096 |  |
| c+ | pr | < | < | 442 | 4.29E-107 |
|  |  |  | >= | 11488 |  |
|  |  | >= | < | 13287 |  |
|  |  |  | >= | 974783 |  |

**Table S2.**
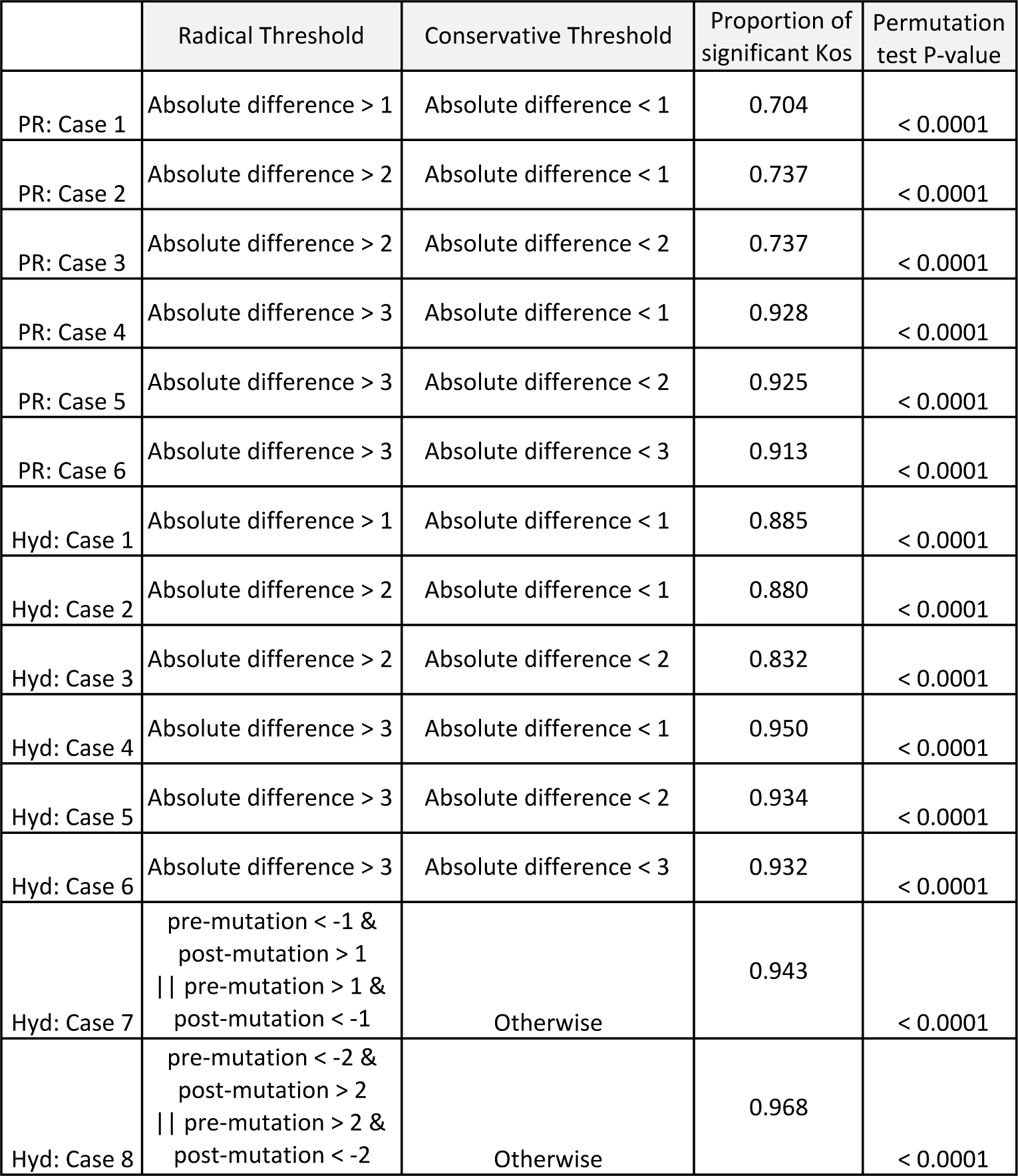
Comparisons between “radical” and “conservative” nonsynonymous substitutions using the polar scale and amino-acid hydropathy index and multiple thresholds for each property. P, permutation test.

